# Identification of potential biomarker candidates of drug-induced vascular injury (DIVI) in rats using gene expression and histopathology data

**DOI:** 10.1101/2022.08.24.505120

**Authors:** Anika Liu, Jordi Munoz-Muriedas, Andreas Bender, Deidre A. Dalmas

**Affiliations:** Centre for Molecular Informatics, Department of Chemistry, University of Cambridge, Lensfield Road, Cambridge, CB2 1EW, UK; Data and Computational Sciences, GlaxoSmithKline, London, UK; GlaxoSmithKline, Non-Clinical Safety, In Vitro In Vivo Translation, Collegeville, PA, USA

**Keywords:** Drug-Induced Vascular Injury, Rat, DIVI, Medial Arterial Necrosis, MAN, Toxicogenomics, Predictive Biomarkers

## Abstract

Drug-induced vascular injury (DIVI) observed in non-clinical species often leads to significant delays or termination of compounds in drug development due to the lack of translatable biomarkers and unknown relevance to humans. This study focused on the identification of potential biomarker candidates of drug-induced vascular injury, or more specifically mesenteric medial arterial necrosis (MAN), in rats. To do so, an adapted bioinformatic filtering pipeline was applied to previously generated gene expression data obtained from laser capture microdissected endothelial and vascular smooth muscle cells and corresponding histopathological annotations from mesenteric arteries following treatment of rats with known vascular toxicants, non-vasotoxic vasoactive comparator compounds, and corresponding vehicle. A novel gene panel including 33 genes with consistent, specific, and dose-responsive dysregulation across multiple treatments inducing MAN was identified. The degree to which these reflect injury progression was characterized and changes were identified in samples from animals where injury was anticipated but not yet histologically observed. The most predictive candidates (AUC > 0.9) with the strongest changes (logFC > 1.7) in MAN encode secreted proteins (*Tnc, Vcan, Timp1* and *Fn1)* which are strongly interlinked with each other and other potential candidate biomarkers encoding cell surface proteins. Although further validation is required for biomarker qualification, the adapted bioinformatic approach utilized in this study provides informed data-driven starting points for further DIVI biomarker discovery and development in rats, as well as potential mechanistic insight into MAN pathogenesis.

## Introduction

Drug-induced vascular injury (DIVI) can be induced within hours of drug administration and is phenotypically identified as morphological changes in the vascular wall including but not limited to medial arterial necrosis (MAN), one of the main hallmark lesions (Kerns et al., 2005). In non-clinical species, DIVI manifests across a wide range of structurally and pharmacologically diverse compound classes (Morton and Houle, 2014). However, the clinical relevance of findings in animals is unclear and several compounds inducing DIVI in model species, such as minoxidil (Sobota, 1989), the phosphodiesterase IV inhibitor apremilast (Kavanaugh et al., 2014) or caffeine (Johansson, 1981), have not been reported to date, to result in DIVI in humans. This poses a hurdle for assessing drug safety, as pre-clinical evidence can result in delays in drug development and termination (Kerns et al., 2005), although it is often unknown whether observations may or may not be relevant to man. One reason for this is the lack of non-invasive, sensitive and specific methods to detect the presence or predict the development of drug-induced vascular lesions, leaving histopathological examination the only option. Caution in this regard arises from a limited understanding of the longer-term consequences of DIVI which may include chronic vascular injury, or cardiovascular morbidity (Pearson et al., 2003).

Mesenteric MAN is predominantly found in medium-sized mesenteric arteries ranging from approximately 100–800 μm, and is hypothesized to be initiated by various mechanisms including hemodynamics effects resulting in biomechanic injury, direct toxicity to vascular cells and/or injury mediated by an immune response (Kerns et al., 2005). This then can lead to damage of vascular tissue with MAN being one of the key hallmark lesions (Dalmas et al., 2011). To support informed decision-making, current research across the industry focuses on identifying non-invasive biomarkers with the ability to detect vascular injury in patients and/or pre-clinical animal models earlier in development and more reliably. Two key consortia with subgroups focusing on research in this area are the Predictive Safety Testing Consortium (PSTC), as part of which the Vascular Injury Working Group (VIWG) studies DIVI in animal models as well as translation across species including humans (https://c-path.org/programs/pstc), and the Innovative Medicines Initiative (IMI) Consortium TransBioLine focusing on DIVI in humans (https://transbioline.com). Both aim to produce evidence towards biomarker identification, development, and qualification and have curated panels of potential circulating protein-based biomarkers (many of which are proprietary) which may reflect mechanisms leading to DIVI pathogenesis based on their association with known histopathological features shared with vascular diseases and/or across species. Those that are described, are related to adaptation in vascular function due to hemodynamic changes, endothelial cell activation and/or inflammatory cell recruitment indicated by endothelial adhesion molecules and pro-inflammatory cytokines, and smooth muscle damage which may lead to leakage of SM-specific proteins, genes or microRNAs into circulation, as well as markers of vascular remodeling which may indicate fibrosis and neovascularization depending on the severity of the histopathological finding. Potential circulating biomarkers noted in literature being evaluated along with their anatomical location by the PSTC VIWG are noted in **Table 1** for ease of reference. Current research across industry consortia including the Predictive Safety Testing Consortium (PSTC) focus efforts on the identification of translatable and predictive biomarkers of DIVI including soluble-based biomarkers which include those related to adaptation in vascular function due to hemodynamic changes, endothelial cell activation and inflammatory cell recruitment indicated by endothelial adhesion molecules and pro-inflammatory cytokines, smooth muscle damage which may lead to leakage of SM-specific proteins into circulation upon necrosis and vascular remodeling which includes fibrosis and neovascularization.

**Table 1:**
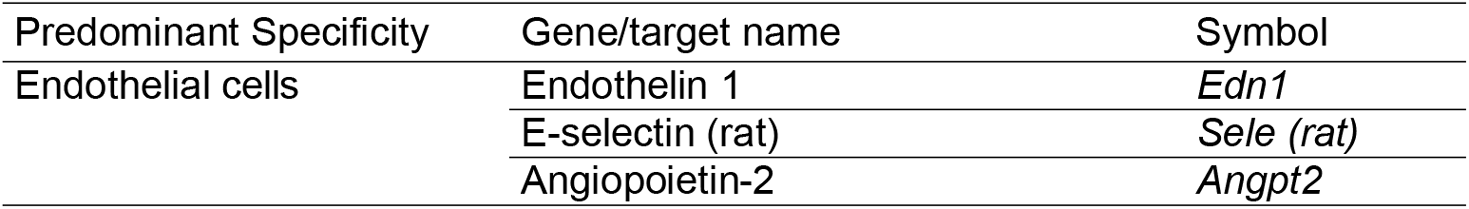
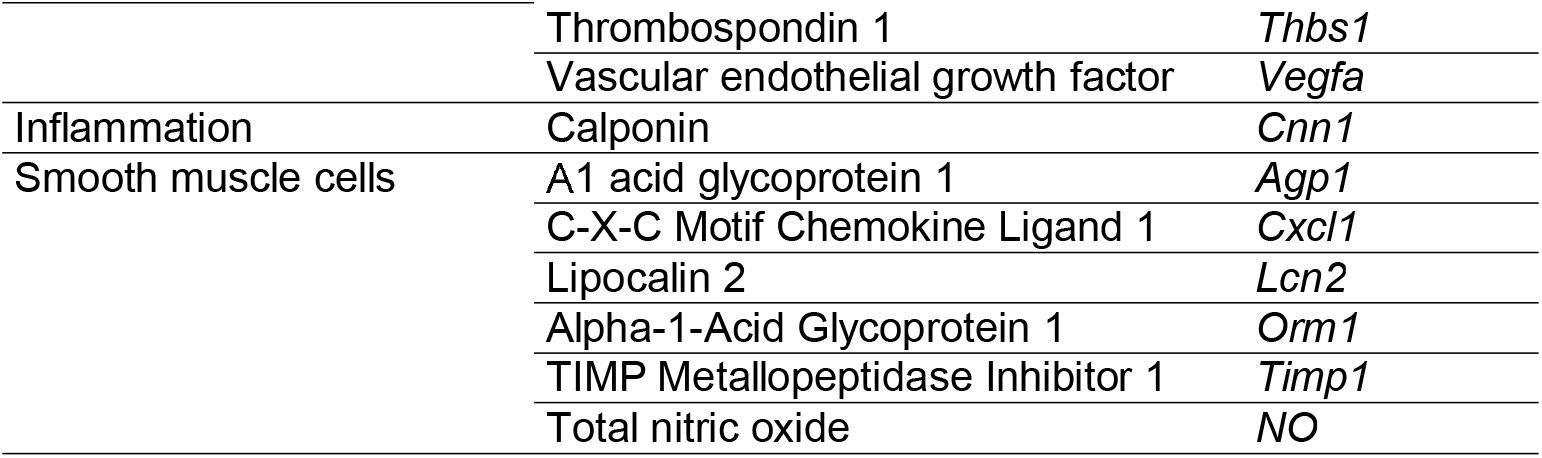
List of potential circulating biomarker candidates prioritized by the Vascular Injury Working Group of the Predictive Safety Testing Consortium (PSTC VIWG) for DIVI qualification and their hypothesized specificity in the prediction of injury pathogenesis. Table adapted from Mikaelian et. al. (2014), with permission from authors.

| Predominant Specificity | Gene/target name | Symbol |
| --- | --- | --- |
| Endothelial cells | Endothelin 1 | *Edn1* |
|  | E-selectin (rat) | *Sele (rat)* |
|  | Angiopoietin-2 | *Angpt2* |
|  | Thrombospondin 1 | *Thbs1* |
|  | Vascular endothelial growth factor | *Vegfa* |
| Inflammation | Calponin | *Cnn1* |
| Smooth muscle cells | A1 acid glycoprotein 1 | *Agp1* |
|  | C-X-C Motif Chemokine Ligand 1 | *Cxcl1* |
|  | Lipocalin 2 | *Lcn2* |
|  | Alpha-1-Acid Glycoprotein 1 | *Orm1* |
|  | TIMP Metalloproteinase Inhibitor 1 | *Timp1* |
|  | Total nitric oxide | *NO* |

Besides efforts to provide additional evidence for potential biomarkers already supported by literature and expert knowledge (Brott et al., 2005; Kerns et al., 2005; Louden et al., 2006a) there have also been investigative efforts to identify additional potential novel candidate biomarkers which are ideally sensitive and specific, mechanistically linked to the pathogenesis and additionally found to precede injury and reflect lesion severity (Louden et al., 2006b; Weaver et al., 2008). In this regard, our prior research (Dalmas et al., 2011) identified candidate tissue biomarkers based on Affymetrix GeneChip analysis from samples of mesenteric arteries of rats treated with multiple vascular toxicants and comparator vasoactive but not vasotoxic compounds (Dalmas et al., 2011), whereas most studies in this area only address few and often single stressors (Daguès et al., 2007a; Dalmas et al., 2008; Heydarkhan-Hagvall et al., 2006; Slim et al., 2003; Weaver et al., 2010, 2008; Zhang et al., 2006) An additional advantage of the data generated in this work, which is also used in this study, is that it is derived from the endothelium and smooth-muscle enriched samples by laser capture microdissection and hence is able to capture changes in the vascular tissue which is not diluted by the neighboring adventitia, connective tissue or lymph nodes. Besides identifying sensitive, specific and dose-responsive potential genomic biomarkers from gene expression data, a subset of the genes was confirmed by quantitative RT PCR (TaqMan) analysis in rats treated for 1 or 4 days with dopamine or fenoldopam where medial arterial necrosis (MAN) was present. For these genes, the absence of changes was further confirmed in rats treated with yohimbine, a vasoactive and non-vasotoxic compound that did not show any evidence of vascular injury in the mesentery (Dalmas et al., 2011, 2008). Furthermore, it was found that some of the potential candidate biomarkers were regulated in mesenteric artery tissue scrape samples from rats treated with fenoldopam approximately 8 hours prior to histological detection of MAN providing further evidence that the genes are good potential candidates for detection of MAN in rats (Dalmas et al., 2011, 2008). However, as previously reported, it is not feasible nor practical to perform this extensive validation by TaqMan across a large set of genes or compounds, and also the biomarker filtering itself might overlook valuable genes by focusing only on a small subset with stringent cut-off criteria (Dalmas et al., 2011).

In this study, a subset of the previously collected microarray gene expression and histopathology data (Dalmas et al., 2011) for selected compounds was further analyzed as part of a collaboration between GSK and the University of Cambridge using an adapted bioinformatic approach to derive an extended set of potential genomic candidate biomarkers which were shown to detect and predict MAN in rats with the aim to provide additional starting points for further DIVI biomarker discovery and development. First, genes were identified with a stronger focus on consistency across treatments at the cost of lower effect size, while still requiring specific, and dose-dependent dysregulation. Further evidence was gained for these genes by investigating the behavior across lesion severity. This approach enabled not only the characterization of genes which correlated with the histological presence of MAN, and injury progression but also the identification of gene expression changes in treatments where MAN is anticipated but not yet microscopically identified. Furthermore, candidate biomarkers were found to encode multiple secreted proteins which may translate to circulatory biomarkers. In addition, an interactive web application was developed using R/Shiny, in which results from this study for genes of interest beyond the 33 genes identified to be most promising candidates for potential biomarker development can be explored interactively. This can be accessed via https://anikaliu.shinyapps.io/divi.and will enable researchers to gather supporting or opposing evidence also for genes of interest which were not shortlisted in this work.

## Methods

### Study design

The data used in this work was provided by GlaxoSmithKline as part of a collaboration with the University of Cambridge. Data was provided from selected compounds from previous experiments (Dalmas et al., 2011), in which male Crl:(CD)SD rats were given various known vasotoxic DIVI-inducing and vaso-active, non-vasotoxic compounds, at three different doses as well as corresponding and vehicle (see Table 2 and Table S 1). Approximately 24 hours following the final dose of each treatment, rats were killed by CO_2_ asphyxiation/exsanguination and necropsied as previously described. The in-life portion of the prior study from which data was obtained was conducted at Charles River Laboratories, Discovery and Development Services (CR-DDS), Argus Division, Horsham, PA, USA. All prior studies were conducted after review by the Charles River Laboratory (Discovery and Development Services, Argus Division, Horsham, PA, USA) Institutional Animal Care and Use Committee (IACUC) in accordance with the GSK Policy on the Care, Welfare and Treatment of Laboratory Animals and were in accordance with the Guide for the Care and Use of Laboratory Animals (NIH Publication, 25, No. 28, 16 August 1996). Endothelium or smooth muscle enriched samples were derived from cryosections of mesenteric arteries through laser capture microdissection, and gene expression levels were measured on Affymetrix Rat Genome 230 2.0 Arrays.

**Table 2:**
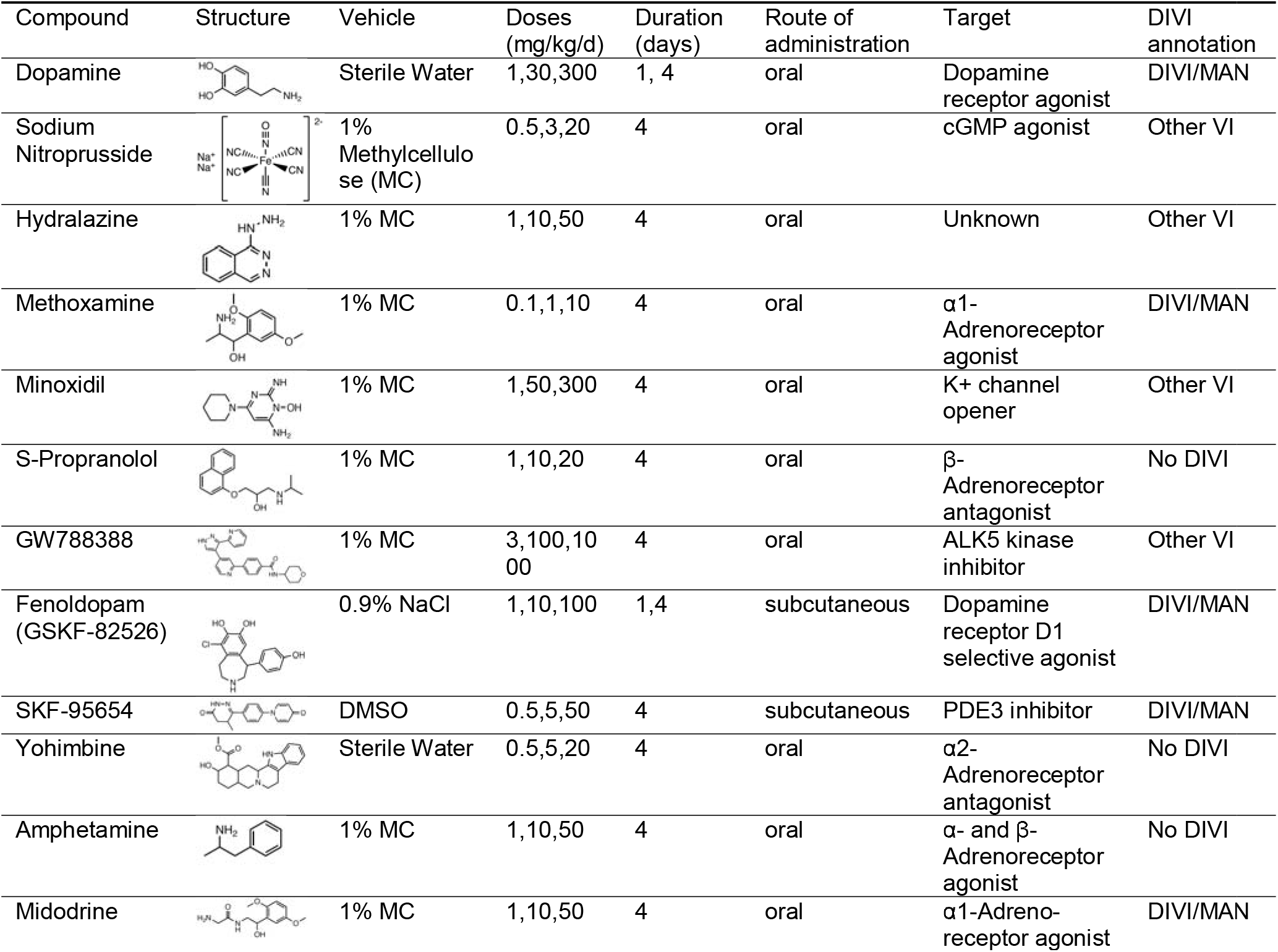
Summary of treatment conditions administered to rats analyzed in this work. The DIVI annotation separates DIVI-inducing compounds which show mesenteric medial arterial necrosis (“DIVI/MAN”) from those which showed other histological changes in the mesenteric artery, such as perivascular and/or fibrinoid necrosis, perivascular fibrosis, EC hypertrophy and/or inflammatory cell infiltration (“Other VI”), and compounds which did not present any of the mentioned histological changes (“No DIVI”). The information was adapted from Dalmas et al. (2011), with permission from the authors.

| Compound | Structure | Vehicle | Doses<br>(mg/kg/d) | Duration<br>(days) | Route of<br>administration | Target | DIVI<br>annotation |
| --- | --- | --- | --- | --- | --- | --- | --- |
| Dopamine |  | Sterile Water | 1,30,300 | 1, 4 | oral | Dopamine<br>receptor agonist | DIVI/MAN |
| Sodium<br>Nitroprusside |  | 1%<br>Methylcellu-<br>lose (MC) | 0.5,3,20 | 4 | oral | cGMP agonist | Other VI |
| Hydralazine |  | 1% MC | 1,10,50 | 4 | oral | Unknown | Other VI |
| Methoxamine | | 1% MC | 0.1,1,10 | 4 | oral | α1-Adrenoreceptor<br>agonist | DIVI/MAN |
| Minoxidil |  | 1% MC | 1,50,300 | 4 | oral | K+ channel<br>opener | Other VI |
| S-Propranolol | | 1% MC | 1,10,20 | 4 | oral | β-<br>Adrenoreceptor<br>antagonist | No DIVI |
| GW788388 |  | 1% MC | 3,100,10<br>00 | 4 | oral | ALK5 kinase<br>inhibitor | Other VI |
| Fenoldopam<br>(GSKF-82526) |  | 0.9% NaCl | 1,10,100 | 1,4 | subcutaneous | Dopamine<br>receptor D1<br>selective agonist | DIVI/MAN |
| SKF-95654 |  | DMSO | 0.5,5,50 | 4 | subcutaneous | PDE3 inhibitor | DIVI/MAN |
| Yohimbine | | Sterile Water | 0.5,5,20 | 4 | oral | α2-Adrenoreceptor<br>antagonist | No DIVI |
| Amphetamine | | 1% MC | 1,10,50 | 4 | oral | α- and β-<br>Adrenoreceptor<br>agonist | No DIVI |
| Midodrine | | 1% MC | 1,10,50 | 4 | oral | α1-Adreno-<br>receptor agonist | DIVI/MAN |

For histopathological analysis, mesentery from each animal was collected, processed, and histopathology was assessed using light microscopy by a single board-certified pathologist, and peer-reviewed by another board-certified pathologist as previously described (Dalmas et al., 2011). Thereby, histopathological findings were documented using severity scores ranging from 0 (absence) to 4 (high severity). Histopathological evidence of MAN (severity score > 0), was identified for 5 out of 12 compounds, including fenoldopam, dopamine, midodrine, methoxamine, and SKF-95654 (see **Table 2**). In contrast, no evidence of vascular injury was observed for yohimbine, S-propranolol, and amphetamine (i.e., no evidence in controls or test-article treated animals of MAN or perivascular and/or fibrinoid necrosis, perivascular fibrosis, endothelial cell hypertrophy or inflammatory cell infiltration).

### Gene expression pre-processing

The raw gene expression levels for treatments evaluated in this study (Table S 1) were first background corrected, log2 transformed, and quantile normalized with RMA (Robust Multi-array Average), accessed through the affy package (Gautier et al., 2004), per study and cell type. Array quality metrics describing distances between array, array intensity distributions and variance mean dependence were computed with the ArrayQualityMetrics package (Kauffmann et al., 2009) and outliers detected based on these statistics were excluded from the downstream analysis. This resulted in overall 304 samples for smooth muscle gene expression, and 300 samples for endothelial gene expression across all treatments. The number of animals in each experimental condition is shown in Table S 1. Study-dependent batch effects were found resulting from differences which are considered to be due mainly to technical variance between studies (each including treatment at multiple doses and its respective vehicle control). Technical variation may have resulted due to multiple experimental factors, such as the age of the rats at the time of treatment, the time of the day at which rats were dosed or necropsied, and differences in the vehicle or and route of administration between compounds. While these batch effects are irrelevant in the case of within-study analysis, which was performed in our prior work (Dalmas et al., 2011), we also analyzed trends across batches in this study. Therefore, batch correction was performed using the ComBat function of the sva package (Leek et al., 2012) regarding the individual studies as batch covariates and using the implemented parametric empirical Bayes framework. The platform information for the Affymetrix Rat Genome 230 2.0 Array was derived from Gene Expression Omnibus (Edgar et al., 2002) using GEO accession GPL1355, and used to aggregate probe IDs to rat gene symbols using the median for all probes uniquely mapping to one gene symbol. Where a probe matched multiple genes, the probe itself was kept avoiding dilution or duplication of the contained information, respectively.

### Filtering of genes

To prioritize genes as potential transcriptomic biomarkers, an adapted approach as compared to the prior work (Dalmas et al., 2011) was used, prioritizing consistency and specificity over effect size in DIVI conditions. To do so, DIVI conditions were first defined as those conditions in which MAN is observed for more than 20% of the animals as a trade-off between including early evidence of DIVI and excluding rare changes (**Table S 1**). The conditions in which MAN was observed are depicted in **Figure 1** showing consistently increasing MAN frequency with increasing dose and MAN across all animals for SKF-95654 and midodrine at the highest dose.

**Figure 1:**
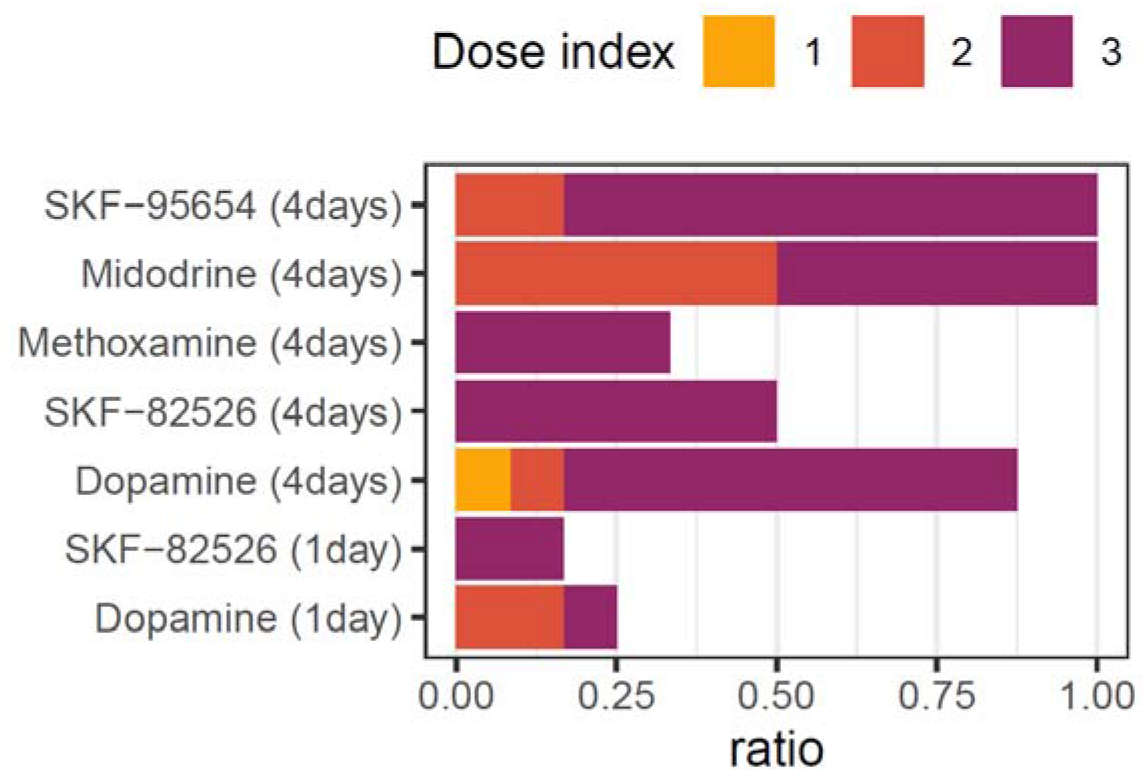
Frequency of Medial Arterial Necrosis (MAN) across conditions. For all conditions in which MAN was found the frequency is shown for each treatment, time and dose. Across all treatments, an increasing frequency of cases is found with increasing dose.

The differential expression for each compound and dose group in comparison to the respective experiment-matched vehicle control was computed using gene-wise linear modeling and empirical Bayes moderated t-statistics provided by the limma R package (Ritchie et al., 2015). Genes which significantly (p-value < 0.05) changed in the same direction across all DIVI conditions (further defined below) with a |logFC| > 0.7 were then identified as conserved genes. While a higher logFC threshold, e.g. |logFC| =1, is more common and was also used in our prior work (Dalmas et al., 2011), this lower logFC cut-off was chosen due to the fact that a gene would be filtered out if a logFC below this threshold was found in any DIVI condition. An alternative approach to balance potential outliers was previously used in our past work (Dalmas et al., 2011) requiring consistency across 8 of the 12 treatments with a |logFC| =1 (**Table S 2**).

Next, genes were removed for which a significant change (p-value < 0.05) in the same direction was also found for any compound at any dose which did not show MAN or other evidence of vascular damage. This includes amphetamine, S-propranolol or yohimbine (**Table S 1**). As potential biomarker genes should reflect the dose-dependent increase in MAN frequency observed across all DIVI compounds, Spearman rank correlation was then computed between gene expression levels of individual animals and the given compound doses, including the vehicle control as a dose of 0 mg/kg, and omitted genes with an absolute Spearman correlation below 0.3 in any of the DIVI conditions. To evaluate whether and how many genes are expected to pass the complete filtering procedure at random we generated a null distribution using 1000 permutations of the compound labels. Only in 0.8% and 1% of permutations, any gene was identified in the endothelium or smooth muscle, respectively, and never were as many genes identified as for the true data indicating that genes are unlikely to pass the filtering procedure by chance (**Figure S 4**).

### Development of an interactive web app

The results using the adapted analyses performed as part of this study with a subset of compounds from our previous work (Dalmas et al, 2011), were used to develop an interactive R/Shinyweb application in order to enable exploration of the results beyond the 33 genes identified as most promising biomarker candidates (https://anikaliu.shinyapps.io/divi). It should be noted, that results on a few genes currently being analyzed internally or as part of other initiatives have been excluded as these data points are currently being analyzed and are intended to be the subject of future publications. This includes but is not limited to data corresponding to the majority of the prioritized candidate biomarkers of the PSTC VIWG which are being analyzed in conjunction with other datasets generated by the consortia as potential supporting evidence for a DIVI non-clinical rat biomarker qualification package.

### Candidate biomarker predictivity

For each gene suggested as a potential candidate biomarker based on this analysis or in previous literature, the ability to predict the presence or absence of MAN was evaluated through the area under the receiver operating characteristic curve (AUC) using the yardstick (Kuhn and Vaughan, 2020) R package. Given that it was in most cases unclear whether an up- or downregulation is expected for literature markers, the higher AUC among both was reported expecting all biomarker candidates to perform better than random.

### Biological annotation

Functional protein-protein associations between conserved genes in both tissues were derived from the STRING database (Szklarczyk et al., 2019) including all interactions with a combined score > 0.4, and the network was visualized in Cytoscape. For gene set analysis, 1,002 gene sets from the Rat Genome Database (Shimoyama et al., 2015) were combined with 2,249 gene sets from MsigDB (Liberzon et al., 2011), canonical pathways (C2 CP) and hallmarks (H) which were mapped from human HGNC symbols to rat gene symbols with BiomaRt (Smedley et al., 2015). In cases where the corresponding rat gene symbol was not represented as an individual gene, it was matched to shared probes if possible. Over-representation of identified genes in these pathway maps was analyzed with the clusterProfiler (Yu et al., 2012) R package using the hypergeometric test statistic and all measured genes as background.

## Results and Discussion

### Identification of transcriptomic biomarker candidates for DIVI

To identify genes with the highest potential for biomarker discovery and development at the cost of losing many other promising genes, an adapted stringent filtering procedure was developed and applied to prioritize genes that show a consistent, specific and dose-dependent response (differences from the previous filtering pipeline are summarized in Table S 2). The number of genes identified at each step is shown in **Figure 2**A. Overall, 33 potential candidate genes with consistent, specific and dose-dependent changes were identified in the smooth muscle- and 10 in the endothelium-enriched samples that correlate with the presence of DIVI. Expression changes for each potential candidate gene across all compounds are shown in **Figure S 2** and **Figure S 3**. For many of the genes, significant changes in expression were frequently identified at lower doses than those for which histopathological evidence of MAN was observed, thus indicating that these changes may precede and predict the occurrence of MAN. This is further supported by the fact that many of the genes identified following 4 daily doses of fenoldopam, one of the conditions in which MAN was observed, were also observed to be regulated 24 hours following a single dose prior to histopathological evidence of MAN.

**Figure 2:**
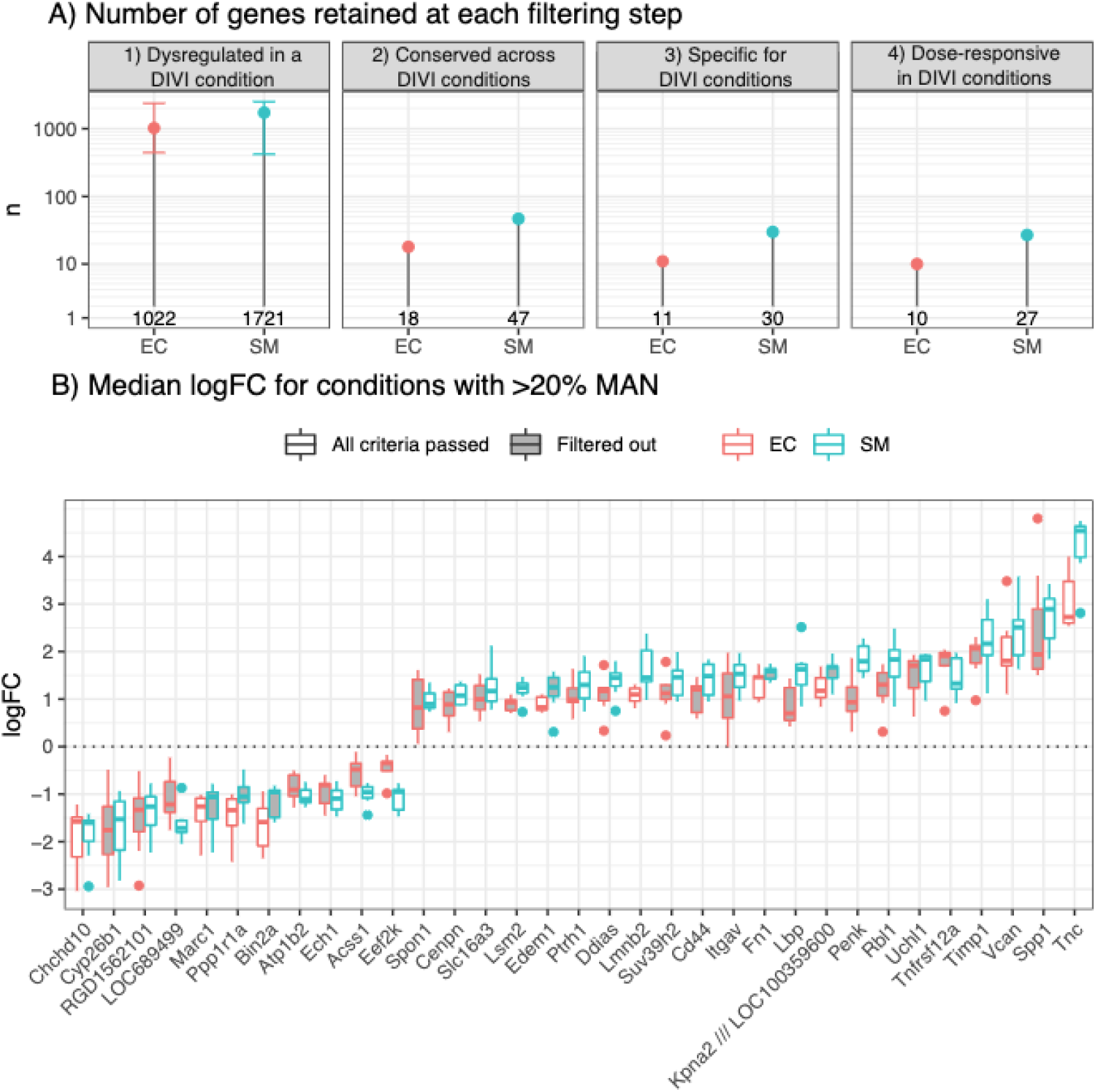
Potential genomic biomarker candidates for MAN identified through filtering criteria. A) Potential gene expression candidate biomarkers for DIVI MAN in endothelial cells (EC) and smooth muscle cells (SM) were identified through multiple filtering criteria using medial arterial necrosis (MAN) as the main readout, and the number of genes is shown for each tissue at each stage. First, differentially expressed genes were identified for each individual condition with >20% MAN. In subsequent steps, only genes with conserved significant differential expression across all DIVI conditions in the same direction, without significant differential expression in any negative control condition and dose-dependent expression changes (|Spearman correlation| ≥ 0.3) were kept. B) Distribution of logFCs for differentially expressed potential candidate genes, identified by the filtering procedure in conditions with >20% MAN.

While all genes passing the filtering criteria show consistent changes across DIVI conditions, the magnitude of expression change varies and is summarized in **Figure 2**B. Overall, the strongest up-regulated gene, as noted by median logFC, in rats with histologic evidence of MAN (i.e. DIVI conditions) in both tissues was found for Tenascin C (*Tnc)*, a gene encoding the glycoprotein Tnc known to be involved in blood vessel injury (Imanaka-Yoshida et al., 2014) which was also identified our prior work while the strongest down-regulation in gene expression was observed for *Chchd10*, encoding mitochondrial Coiled-coil-helix-coiled-coil-helix domain-containing protein 10 (**Figure 2**B) highlighting that these might be easily detectable. For all potential candidate genes identified, consistent directionality of dysregulation in both tissues was observed.

The list of candidate genes identified in this study was then compared to the ones from the previous work (Dalmas et al., 2011), and 6 out of 33 biomarker candidates were found to overlap with the previously proposed 57 genes (Table S 4). Out of the six genes that were noted to be similarly regulated in a dose-responsive manner across the analysis pipelines, Tissue inhibitor of Metallopeptidase 1 (*Timp1)*, Fibronectin 1 (*Fn1*), Karyopherin Subunit Alpha 2 (*Kpna2*) and Versican (*Vcan*) have been previously confirmed to be regulated using quantitative RT-PCR (TaqMan), as further outlined in Table S 4. While none of the results were disproved, results for Tenascin C (*Tnc*) were undetermined for fenoldopam and undetermined across all treatments for Peptidyl-TRNA Hydrolase 1 Homolog (*Ptrh1*). Furthermore, three of these genes were already prioritized by the PSTC VIWG (**Table 1**), namely *Timp1*, which was first suggested as a potential vascular injury biomarker by Dagues et al. (Daguès et al., 2007b), *Fn1* and *Tnc*. Thus, the adapted analysis workflow is able to recover previously confirmed results, as well as able to identify and prioritizes new genes as potential candidate biomarkers.

The differences in the number of genes identified between the current and prior work (Dalmas et al, 2011) can partially be explained by differences in filtering criteria (**Table S 2**), as well as the fact that analyses in this study were performed on a subset of compounds from past work (Dalmas et al, 2011). On a more general level, both filtering pipelines aimed at prioritizing a short list of most promising genes and do not claim that other genes are not unsuitable as biomarker candidates. If new biomarker candidates are proposed in the future, it may be of interest to understand across which DIVI conditions expression changes, indicative of MAN, are observed. Hence, a R/Shiny web application was developed using which the behavior of a gene of interest can be inspected without requiring any background in bioinformatics or programming (https://anikaliu.shinyapps.io/divi). In the application, the differential expression results across all treatments can be visualized as well as other metrics which were computed in this study.

### Characterization of biomarker candidate gene properties

To evaluate the ability of the potential biomarker candidate expression levels to separate animals with and without MAN, the area under the ROC curve (AUC) was computed for each candidate biomarker prioritized in this study and previous work (Dalmas et al., 2011). This further ranks the individual genes with respect to their predictivity observed in the current dataset and overall identifies correlated performance in both tissues indicating that the derived performance should be representable for the vascular tissue (Figure 3). Among the genes identified as potential biomarker candidates for the prediction of MAN in rats in this study, the highest AUCs are observed for *Fn1* in the smooth muscle (AUC of 0.97) and *Timp1* in the endothelium (0.94), while the lowest AUCs are observed for *Bin2a* (0.81) and *Eef2k* (0.73) in smooth muscle and endothelium, respectively. As expected, there was an overall high predictive performance observed in both endothelium (AUC between 0.59-0.94) and smooth muscle (AUC between 0.56-0.97) also for the genes previously identified in the prior study (Dalmas et al., 2011). In this regard, *Cbx7* should be highlighted which reaches an even higher performance in the endothelium (0.94) than genes prioritized in this study as well as high performance in the smooth muscle (0.95). However, it did not show sufficiently strong dysregulation across DIVI conditions and was hence not included. In contrast, the lowest performance was found for ADAM Metallopeptidase Domain 9 (Adam9) with an AUC of 0.60 in the endothelium and 0.56 in the smooth muscle.

**Figure 3:**
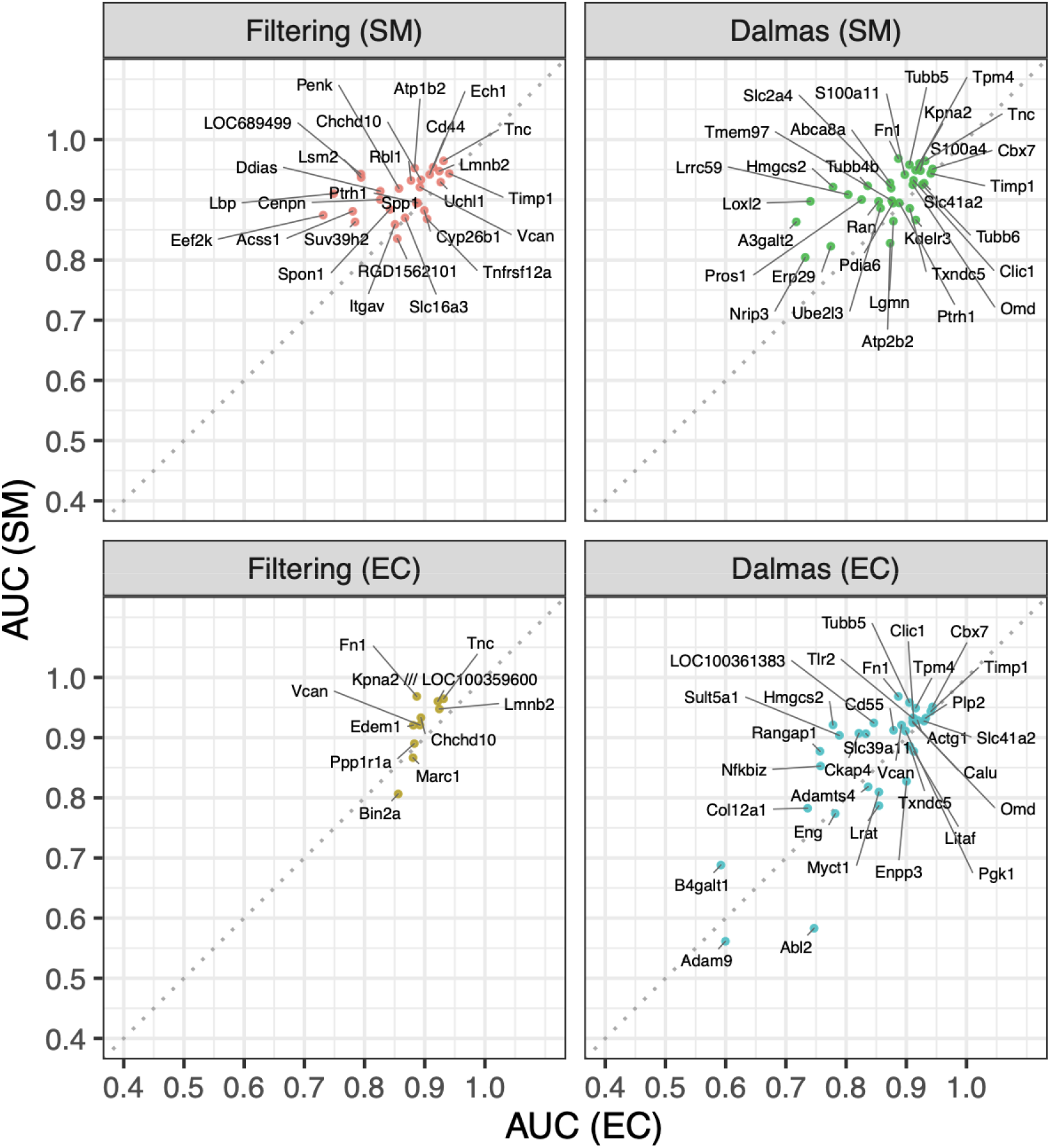
Ability of potential biomarker candidates to separate individuals with and without MAN. For each potential biomarker candidate gene, the ability to separate animals with and without evidence of medial arterial necrosis (MAN) based on expression in the endothelium (EC) or smooth muscle (SM) is shown as AUC. Overall, genes identified in this study and prior work (Dalmas et al., 2011) were found to achieve high AUCs.

The association between expression change and lesion severity was then analyzed to identify genes which reflect injury progression and show detectable expression changes prior to or during early pathogenesis, as these are desired properties for safety biomarkers (Brott et al., 2005; Louden et al., 2006b) (**Figure 1, Table S 1**). Therefore, groups of animals were defined based on the observed severity score for MAN and the expression levels in these samples were compared to those from animals only treated with corresponding vehicle control. Animals without evidence of MAN (severity score of 0) were additionally separated into animals treated with corresponding negative control which should not show changes in marker expression if these genes are predictive of MAN in rats, and animals in DIVI conditions which may show changes in gene expression despite the absence of morphological changes, as well as other conditions. The results of this analysis are shown in **Figure 4** and strong and largely significant changes were identified across lesions of all severities as well as animals in DIVI conditions without MAN, while significant changes are not observed for negative control or other DIVI unrelated treatments. This indicates that changes on the transcriptomic level for selected genes might not only correlate with the presence of injury but can potentially also be predictive of impending vascular damage. While some potential candidate genes remain largely constant across all lesion severities and animals in DIVI conditions without MAN, such as Lamin B2 (*Lmnb2)* or Beta-galactosidase *(Bin2a)*, others show an increasing change with increasing severity scores potentially reflecting vascular injury progression, e.g., ER degradation-enhancing alpha-mannosidase-like protein 1 (*Edem1), Timp1*, TNF Receptor Superfamily Member 12A (*Tnfrsf12a)* or Cytochrome P450 Family 26 Subfamily B Member 1 (*Cyp26b1)*. However, comparisons across lesion severity should be treated with caution due to the low number of animals found with severe MAN indicated by a severity score of 3, and more generally due to the fact that this analysis was specific for MAN and other vascular changes were not included (**Table S 3**).

**Figure 4:**
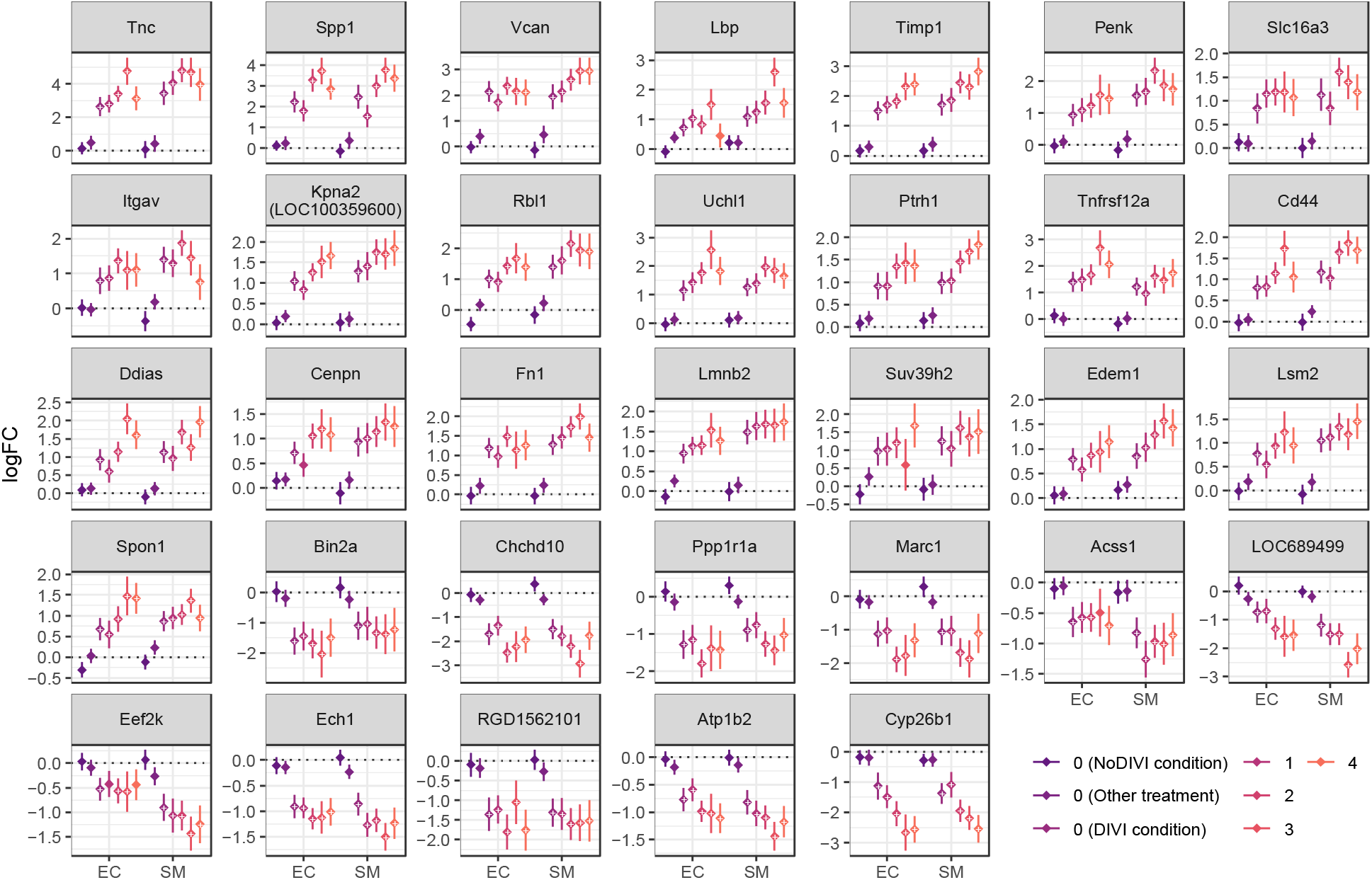
Gene expression changes in medial arterial necrosis (MAN) severity groups in comparison to vehicle controls. Animals which received a treatment were separated into groups based on the observed presence specifically of histopathology. Animals with MAN were grouped by the observed severity, and animals without MAN were divided based on the respective experimental condition: Negative control treatments (“NoDIVI condition”), conditions with >20% MAN (“DIVI condition”) and others which showed morphological changes in the vasculature but not MAN (“Other treatments”). For each group, significant differential expression in comparison to vehicle control (FDR < 0.05) is indicated by “+” and the logFC incl. 95% confidence interval is shown. This identifies significant dysregulation across all genes for animals in DIVI conditions, even without or potentially prior to histologic evidence of MAN. Furthermore, severity-dependent differential expression magnitude is identified for some genes, including *Timp1* and *Cyp26b1*.

### Biological context of biomarker candidate genes

To better understand the potential biological role of the identified genes, protein-protein interactions between proteins encoded by genes with conserved dysregulation across DIVI conditions in both tissues were derived. Overall, more interactions than expected at random (PPI enrichment p-value < 10^−16^) were identified, as well as two clusters of functionally associated genes (**Figure 5**). The bigger cluster includes largely ECM-related proteins including extracellular proteins Fn1, Timp1, Tnc and Vcan which are detected in both tissues and central nodes in the cluster. Cellular receptors in the cluster linked to those proteins include the integrins alpha V (Itgav) and alpha 5 (Itga5), which interact with Fn1 and are involved in fibronectin matrix formation and cardiovascular development (Chen et al., 2015).

**Figure 5:**
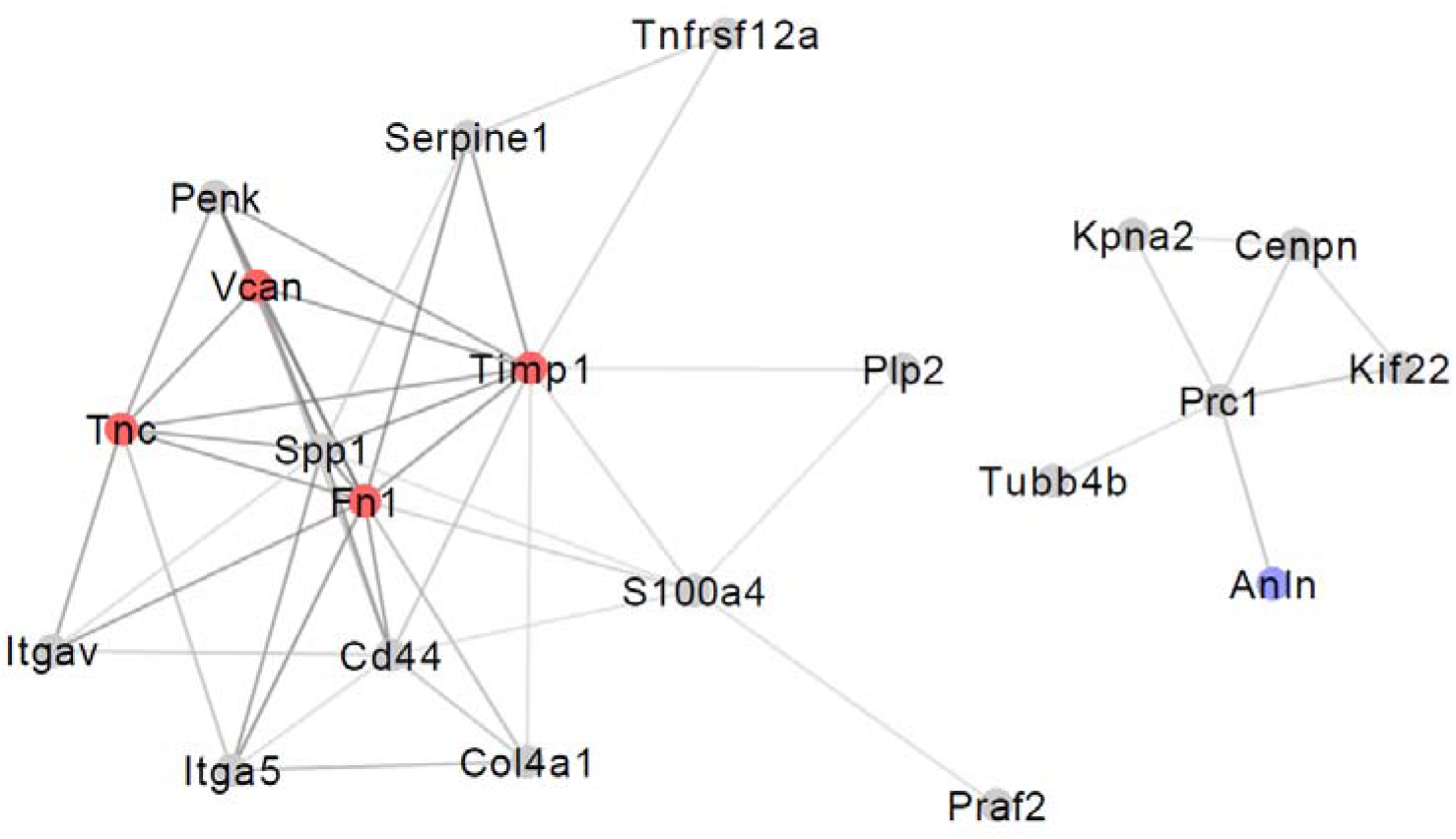
Functional protein-protein associations between conserved genes. Interactions from the STRING database with medium confidence (confidence score>0.4) between conserved genes were derived. A greater number of associations are found than expected at random (13 interactions expected, PPI enrichment p-value < 10-16). Genes identified in both tissues (labeled in red) are found at the core of the bigger cluster. Only one of the genes identified in the endothelium, Anln labeled in blue, showed functional associations, while the majority of genes and interactions are found in the smooth muscle (grey).

Furthermore, secreted phosphoprotein 1 (Spp1) was identified which is increased in multiple vascular diseases (Lok and Lyle, 2019) and is known to interact with both reported integrins as well as the hyaluronic acid receptor Cd44. The second cluster contains tubulin beta-4B chain (Tubb4b) and the Kinesin-Like Protein (Kif22), which are related to microtubule-based transport, Anillin (Anln) which is involved in cytokinesis, the centromere protein N (Cenpn) and Kpna2. All of which are linked to the protein regulator of cytokinesis 1 (Prc1), the central node in this cluster, pointing to vascular hyperplasia. Hence, plausible associations between proteins encoded by candidate genes were identified shedding further light on their potential mechanistic role.

An over-representation analysis was then performed for each tissue using conserved genes or only biomarker candidate genes, respectively. From this analysis, three pathways in the endothelium and twenty-two in the smooth muscle were identified, which can be explained by the higher number of genes observed in the smooth muscle (**Figure 6**). In both tissues, epithelial-mesenchymal transition (EMT) is over-represented which is a key process in tissue repair and fibrosis during which epithelial cells switch to a mesenchymal phenotype which is, amongst others, characterized by the expression of *Fn1* and shows strongly modulated interactions with the ECM (Liberzon et al., 2015). While epithelial cells are not a key component in the vasculature, EMT is closely related to endothelial-mesenchymal transition (EndMT) which has been previously observed in various cardiovascular diseases (Kovacic et al., 2019). A subset of the genes involved in EMT is additionally linked to ECM proteoglycans which are the second gene set overrepresented in both tissues. These are generally known to be upregulated in early vascular lesions and take in multiple key roles in vascular injury, such as mechanotransduction, regulation of leukocyte invasion and inflammation, control over blot clotting, ECM organization as well as vascular calcification (Fu and Tarbell, 2013). The third shared over-represented pathway is related to the ECM and is termed “Regulation of Insulin-like Growth Factor (IGF) transport and uptake by Insulin-like Growth Factor Binding Proteins (IGFBPs)”. However, to note, the identified extracellular proteins in this pathway are not directly related to IGF, but phosphorylated by the same kinase as multiple IGFBPs, namely the extracellular serine/threonine protein kinase FAM20C, which is responsible for the majority of extracellular phosphorylations (Tagliabracci et al., 2015). In the smooth muscle, multiple integrin-related pathways, such as focal adhesions, were identified which are related to the two RGD-motif binding integrins integrin-alpha-V (Itgav) and integrin-alpha-5 (Itga5), as well as the extracellular proteins Col4a1 (Arresten), Fn1, Tnc and Spp1 which interact with integrins on the membrane surface. Additionally, Cd44 is included in the “Integrin cell surface interactions” gene set, potentially due to the known crosstalk with Osteopontin (Spp1)-induced signaling described in the Avb3 OPN pathway. Additional extracellular proteins classified as core matrisome by Naba et al. (Naba et al., 2012) are Spondin 1 (Spon1) and Osteomodulin (Omd). Also, cell surface interactions at the vascular wall are identified, which indicate leukocyte extravasation, and are linked to the two integrins and their interaction partners Fn1 and Cd44 as well as the surface proteins Sodium/potassium-transporting ATPase subunit beta-2 (Atp1b2) and the Monocarboxylate transporter 4 (Slc16a3) which both interact with the extracellular matrix metalloproteinase inducer Basigin. The over-represented parent process hemostasis additionally includes Tubb4b, Kif22 and the plasminogen activator inhibitor 1 (Serpine1), which is involved in the controlled degradation of blood clots via the Urokinase-type plasminogen activator (uPA) and uPAR-mediated signaling. This analysis hence provides further insight into the potential mechanistic interplay between the identified conserved and potential candidate marker genes and indicates potential biological pathways leading to the development and progression of MAN. In particular, changes on the cell surface and interaction with the ECM are highlighted, which are known processes involved in vascular injury and remodeling (Ponticos and Smith, 2014).

**Figure 6:**
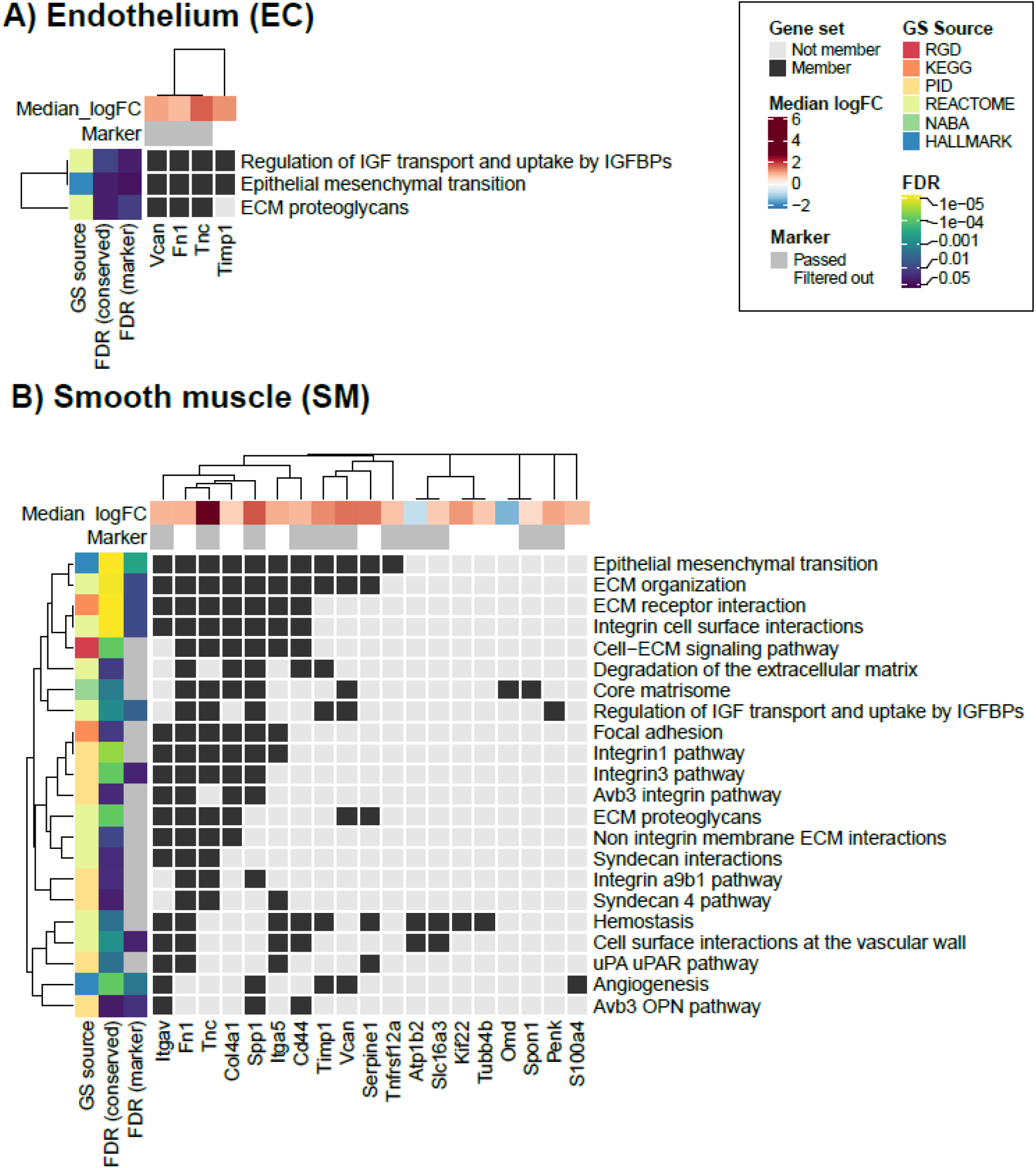
Enriched pathways in conserved and potential marker genes. The gene set membership is shown for conserved genes and significantly enriched pathways (FDR<0.05) in endothelium (A) and smooth muscle (B), respectively. The median logFC across conditions with >20% MAN is shown for all genes and marker genes are additionally highlighted. For the genesets, FDR among conserved and marker genes is shown to identify genesets only enriched among conserved genes. Moreover, the geneset source is annotated with all genesets except the ones from Rat Genome Database being derived through MSigDB.

## Conclusion

In this study, potential genomic biomarker candidates for MAN, a histological indicator of DIVI, in rats were identified using Affymetrix GeneChip data obtained from smooth muscle- and endothelium-enriched mesenteric artery samples and corresponding histopathology data from rats treated with selected compounds evaluated in previous work, i.e. 12 compounds including vasotoxic compounds known to elicit MAN and vasoactive non-vasotoxic comparator compounds at multiple doses including vehicle controls (Dalmas et al., 2011). An adapted bioinformatic gene filtering pipeline, prioritizing consistency and specificity over a larger effect size was used in this study in comparison to our previous work. In both studies, the expression of the candidate gene was required to be dose-responsive as a proxy for whether the genes reflect increasing injury. In this study, gene expression changes across lesion severity were also characterized as a surrogate for lesion progression and revealed that animals in DIVI conditions which do not yet show histopathological evidence of MAN show changes in expression suggesting that the identified biomarker candidates might predict the occurrence of DIVI, specifically MAN.

From a biological perspective, genes encoding proteins involved in ECM interactions were identified to be significantly enriched among the shortlisted candidate genes, and also *Tnc, Timp1, Spp1* and *Fn1* encoding secreted proteins were found to show the highest up-regulation and predictivity. Additionally, it was identified that these genes, as well as the integrins *Itgav* and *Itga5*, which are interaction partners on the cell surface, are highly connected through protein-protein associations further indicating a joint mechanistic role. It should also be highlighted that genes which encode secreted proteins have higher chances of translating to circulating biomarkers (Kerns et al., 2005) and could hence potentially be detected directly from plasma or serum in a non-invasive manner.

Overall, this work not only provides an extended list of promising potential candidate genomic biomarkers for the identification of MAN in rats based on sensitivity, specificity and dose-response but also provides additional supporting evidence by analyzing the newly identified genes’ abilities to reflect lesion severity and the potential mechanistic role in MAN pathogenesis. Given the continued unmet need for DIVI biomarkers, the potential genomic biomarker candidates for MAN and DIVI identified in this work provide valuable data-driven starting points for biomarker discovery and additional follow-up investigations. While only results for the most promising candidate biomarkers are shown, it is clear that biomarker development in practice is heavily informed by broader knowledge on DIVI as demonstrated by ongoing work of the PSTC VIWG (https://c-path.org/programs/pstc) and TransBioLine (https://transbioline.com). To also support future efforts, this work also provides a publicly available application allowing visualization and exploration of gene-level results associated with the presence and/or absence of MAN in rat mesentery following treatment with known vasotoxic as well as vaso-active but non-vasotoxic compounds in comparison to corresponding vehicle controls.

Although additional follow-up work is needed, including confirmation of gene expression changes in tissue from mesentery samples of rats with MAN as well as further investigation of injury initiation and progression using time course studies, the results in this study offer potential new avenues to investigate potential translatable biomarkers of MAN and DIVI in rats.

## Conflict of Interest

AL is funded by GSK and DA is an employee at GSK. JM was an employee at GSK and is now an employee at UCB.

## Author Contributions

***Anika Liu:*** Conceptualization, Methodology, Software, Formal analysis, Investigation, Data curation, Visualization, Writing – original draft; ***Jordi Munoz-Muriedas:*** Funding Acquisition, Project Administration, Supervision, Resources, Validation, Data curation, Writing – review and editing; ***Andreas Bender:*** Funding acquisition, Project administration, Supervision, Resources, Writing – review and editing; ***Deidre A. Dalmas:*** Conceptualization, Resources, Validation, Investigation, Supervision, Data Curation, Formal Analysis, Writing –review and editing.

## Acknowledgments

The authors would like to thank Randall Smith for logistical support on this study as well as the team which was involved in the original study and data generation including Heath Thomas, and Angela-Hughes Earle for histopathological evaluation and/or direction of the study design and support for the in vivo studies from which the data was obtained. The authors also thank Janice Kane, David Mullins, Tushar Chordia, Matthew Steinga, Pratik Patel, Yifen Chen, and Georgina Paulazzo for the preparation and processing of RNA samples and/or Affymetrix GeneChip Analysis and Molly Cool-Bainter for in-life logistical study support from which the originating samples were generated.

## Figure Legends

**Figure S 1: Effect of logFC cutoff on the number of genes retained during the filtering procedure**

**Figure S 2: Expression distribution across experiments for markers identified in the endothelium.**

**Figure S 3: Expression distribution across experiments for markers identified in the smooth muscle.**

**Figure S 4: Number of genes identified in filtering with permutated labels.**

**Figure S 5: Differential expression in the endothelium for previously proposed DIVI biomarker candidates (Dalmas et al., 2011).**

**Figure S 6: Differential expression in the smooth muscle for previously proposed DIVI biomarker candidates (Dalmas et al., 2011).**

**Table S 1:** Frequency of mesenteric arterial necrosis (MAN) across treatments. For each treatment and dose used on this study, the number of animals with MAN is shown followed by the number of animals without MAN. As in some cases, gene expression in the endothelium or smooth muscle was not available or removed in the quality control, the number of samples for which data were available from each is shown.

| Treatment | Histopathology |  |  |  | Endothelium |  |  |  | Smooth muscle |  |  |  |
| --- | --- | --- | --- | --- | --- | --- | --- | --- | --- | --- | --- | --- |
|  | Presence of MAN |  |  |  | Number of samples available for analysis from animals with MAN |  |  |  |  |  |  |  |
| Dose index | 0 | 1 | 2 | 3 | 0 | 1 | 2 | 3 | 0 | 1 | 2 | 3 |
| Dopamine (1 day) | 0\|5 | 0\|6 | 1\|5 | 1\|3 | 0\|5 | 0\|6 | 1\|5 | 1\|3 | 0\|5 | 0\|6 | 1\|5 | 1\|3 |
| SKF-82526 (1 day) | 0\|10 | 0\|11 | 0\|11 | 2\|10 | 0\|10 | 0\|8 | 0\|9 | 2\|9 | 0\|10 | 0\|10 | 0\|11 | 1\|10 |
| Dopamine (4 days) | 0\|10 | 1\|11 | 2\|10 | 7\|1 | 0\|10 | 1\|10 | 2\|9 | 4\|1 | 0\|9 | 0\|10 | 2\|7 | 6\|1 |
| SKF-82526 (4 days) | 0\|9 | 0\|9 | 0\|8 | 4\|4 | 0\|9 | 0\|8 | 0\|8 | 4\|3 | 0\|9 | 0\|9 | 0\|8 | 3\|4 |
| Methoxamine (4 days) | 0\|6 | 0\|6 | 0\|6 | 2\|4 | 0\|5 | 0\|6 | 0\|5 | 2\|4 | 0\|6 | 0\|5 | 0\|4 | 2\|4 |
| Midodrine (4 days) | 0\|6 | 0\|6 | 3\|3 | 4\|0 | 0\|5 | 0\|5 | 3\|3 | 4\|0 | 0\|5 | 0\|6 | 3\|3 | 4\|0 |
| SKF-95654 (4 days) | 0\|6 | 0\|6 | 1\|5 | 5\|0 | 0\|6 | 0\|5 | 1\|5 | 4\|0 | 0\|5 | 0\|6 | 1\|5 | 5\|0 |
| Amphetamine (4 days) | 0\|6 | 0\|6 | 0\|6 | 0\|0 | 0\|5 | 0\|6 | 0\|6 | 0\|0 | 0\|5 | 0\|6 | 0\|6 | 0\|0 |
| S-Propranolol (4 days) | 0\|6 | 0\|5 | 0\|6 | 0\|6 | 0\|5 | 0\|3 | 0\|5 | 0\|6 | 0\|6 | 0\|5 | 0\|5 | 0\|4 |
| Yohimbine (4 days) | 0\|6 | 0\|6 | 0\|6 | 0\|6 | 0\|5 | 0\|6 | 0\|6 | 0\|4 | 0\|6 | 0\|5 | 0\|5 | 0\|6 |
| Hydralazine (4 days) | 0\|5 | 0\|5 | 0\|6 | 0\|6 | 0\|5 | 0\|5 | 0\|6 | 0\|6 | 0\|5 | 0\|5 | 0\|6 | 0\|6 |
| Sodium Nitroprusside (4 days) | 0\|5 | 0\|5 | 0\|6 | 0\|5 | 0\|5 | 0\|5 | 0\|6 | 0\|5 | 0\|5 | 0\|5 | 0\|6 | 0\|5 |
| Minoxidil (4 days) | 0\|6 | 0\|6 | 0\|6 | 0\|5 | 0\|5 | 0\|5 | 0\|6 | 0\|5 | 0\|6 | 0\|5 | 0\|6 | 0\|4 |
| GW788388 (4 days) | 0\|6 | 0\|6 | 0\|6 | 0\|6 | 0\|5 | 0\|5 | 0\|6 | 0\|6 | 0\|6 | 0\|6 | 0\|4 | 0\|6 |

**Table S 2:**
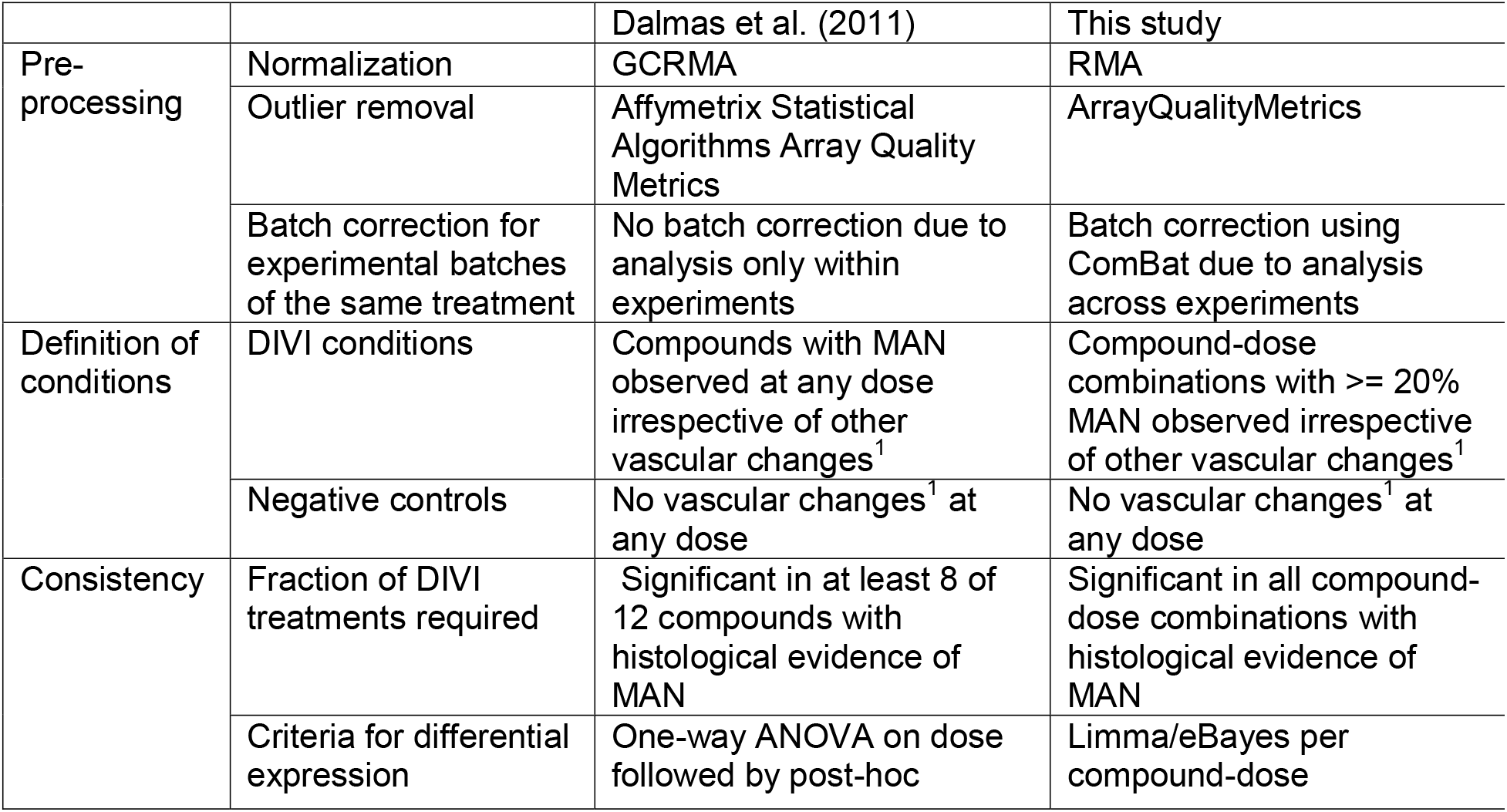
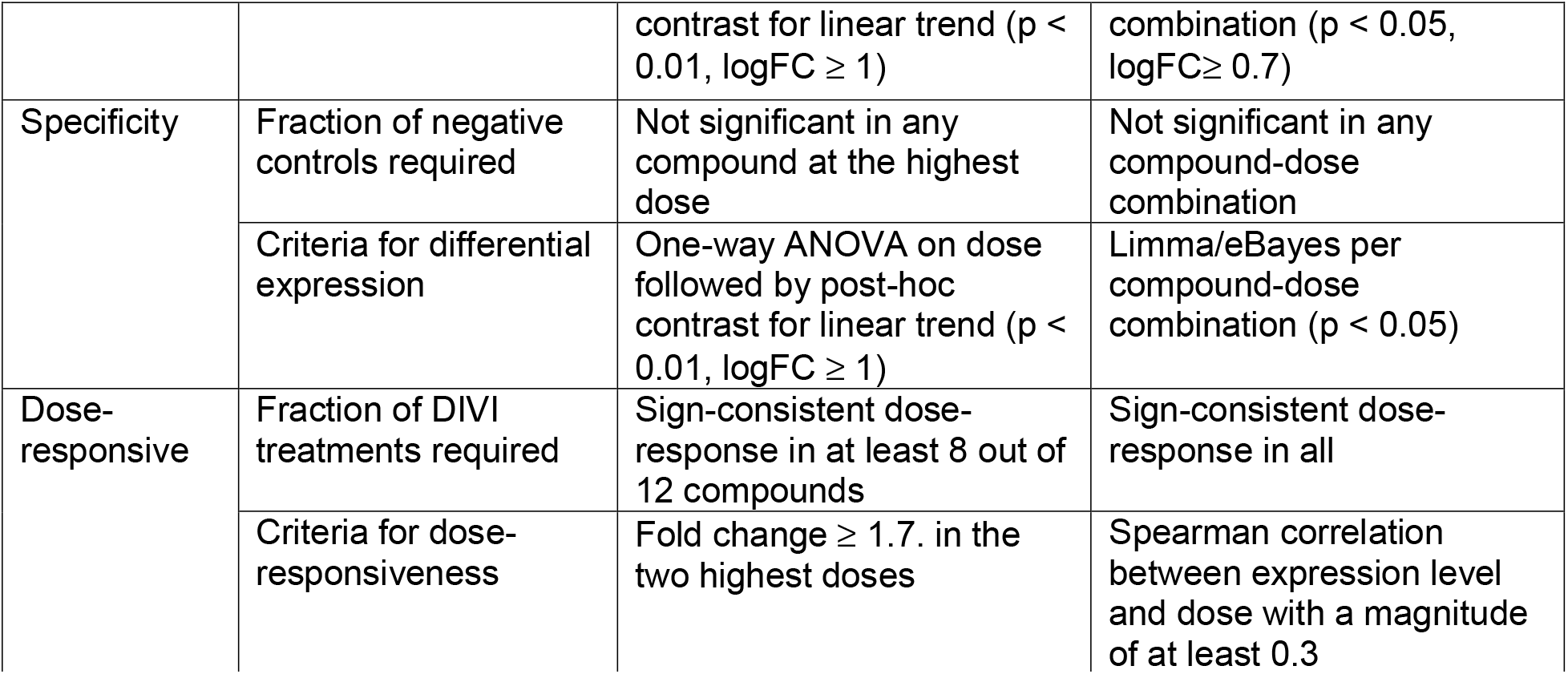
Comparison between filtering implemented by Dalmas et al. (2011) and this study. Vascular changes are defined as mesenteric medial arterial necrosis, perivascular and/or fibrinoid necrosis, perivascular fibrosis, EC hypertrophy and/or inflammatory cell infiltration.

**Table S 3:** Number of samples by MAN group.

| MAN group | EC | SM |
| --- | --- | --- |
| 0 (Control) | 85 | 88 |
| 0 (NoDIVI condition) | 42 | 42 |
| 0 (Other treatment) | 164 | 166 |
| 0 (DIVI condition) | 14 | 15 |
| 1 | 10 | 9 |
| 2 | 11 | 9 |
| 3 | 3 | 6 |
| 4 | 5 | 5 |

**Table S 4:** Genes with conserved and specific dysregulation in DIVI. The identified genes in smooth muscle (SM) and endothelial cells (EC) are shown, including information on whether this was also reported in our prior work (Dalmas et al., 2011) or prioritized by VIWG (Mikaelian et al., 2014). Furthermore, previous quantitative RT-PCR (TaqMan) results are indicated in which expression in enriched individual tissue types and/or tissue scrapes from the entire mesentery of rats treated for 4 days with dopamine (300 mg/kg/day) and fenoldopam (100 mg/kg/day), as well as of rats treated with yohimbine (20 mg/kg/day) as a negative control, was measured.

| Gene Symbol | Gene name | Tissue | Literature | TaqMAN |
| --- | --- | --- | --- | --- |
| Up-regulated genes |  |  |  |  |
| Cd44 | Cd44 molecule | SM | - | - |
| Cenpn | centromere protein N | SM | - | - |
| Ddias | DNA damage-induced apoptosis suppressor | SM | - | - |
| Edem1 | ER degradation enhancer, mannosidase alpha-like 1 | EC | - | - |
| Fn1 | fibronectin 1 | EC | Dalmas (SM\|EC) | Up |
| Itgav | integrin, alpha V | SM | - | - |
| Kpna2 | karyopherin alpha 2 /// | EC | Dalmas | Up |
| (LOC100359600) | karyopherin alpha 2-like |  | (SM)\|VIWG |  |
| Lbp | lipopolysaccharide binding protein | SM | - | - |
| Lmnb2 | lamin B2 | SM\|EC | - | - |
| Lsm2 | LSM2 homolog, U6 small nuclear RNA associated (S. cerevisiae) | SM | - | - |
| Penk | proenkephalin | SM | - | - |
| Pthr1 | peptidyl-tRNA hydrolase 1 homolog (S. cerevisiae) | SM | Dalmas (SM) | Ct values undetermined for all samples |
| Rbl1 | retinoblastoma-like 1 (p107) | SM | - | - |
| Slc16a3 | solute carrier family 16 (monocarboxylate transporter), member 3 | SM | - | - |
| Spon1 | spondin 1, extracellular matrix protein | SM | - | - |
| Spp1 | secreted phosphoprotein 1 | SM | - | - |
| Suv39h2 | suppressor of variegation 3-9 homolog 2 (Drosophila) | SM | - | - |
| Timp1 | TIMP metalloproteinase inhibitor 1 | SM | Dalmas (SM\|EC)\|VIWG\|SAFE-T | Up |
| Tnc | tenascin C | SM\|EC | Dalmas (SM) | Partial (Ct values undetermined for Fenoldopam) |
| Tnfrsf12a | tumor necrosis factor receptor superfamily, member 12a | SM | - | - |
| Uchl1 | ubiquitin carboxyl-terminal esterase L1 (ubiquitin thiolesterase) | SM | - | - |
| Vcan | versican | SM\|EC | Dalmas (EC) | Up |
| Down-regulated genes |  |  |  |  |
| Acss1 | acyl-CoA synthetase short-chain family member 1 | SM | - | - |
| Atp1b2 | ATPase, Na<sup>+</sup>/K<sup>+</sup> transporting, beta 2 polypeptide | SM | - | - |
| Chchd10 | coiled-coil-helix-coiled-coil-helix domain containing 10 | SM\|EC | - | - |
| Cyp26b1 | cytochrome P450, family 26, subfamily b, polypeptide 1 | SM | - | - |
| Ech1 | enoyl CoA hydratase 1, peroxisomal | SM | - | - |
| Marc1 | mitochondrial amidoxime reducing component 1 | EC | - | - |
| Ppp1r1a | protein phosphatase 1, regulatory (inhibitor) subunit 1A | EC | - | - |
| RGD1562101 | similar to very large G-protein coupled receptor 1 | SM | - | - |
| Eef2k | eukaryotic elongation factor-2 kinase | SM | - | - |
| Bin2a | beta-galactosidase-like protein | EC | - | - |
| LOC689499 | similar to Y97E10AL.1 | SM | - | - |

**Table S 5:** Functional protein-protein interactions between conserved proteins derived from STRING. The STRING scores for all observed sources of evidence are shown for all associations with at least middle confidence requiring a combined probability that the interaction is true, or short combined score, above 0.4. The two proteins, Protein 1 and Protein 2, are thereby provided in alphabetical order.

| Protein 1 | Protein 2 | Homo-logy | Co-expression | Experiment | Database | Text-mining | Combined score |
| --- | --- | --- | --- | --- | --- | --- | --- |
| Cd44 | Spp1 | 0 | 0.088 | 0.181 | 0.9 | 0.857 | 0.987 |
| Fn1 | Itga5 | 0 | 0.079 | 0.494 | 0.9 | 0.763 | 0.987 |
| Fn1 | Serpine1 | 0 | 0.12 | 0.055 | 0.9 | 0.79 | 0.98 |
| Spp1 | Timp1 | 0 | 0.257 | 0 | 0.9 | 0.757 | 0.98 |
| Fn1 | Spp1 | 0 | 0.064 | 0 | 0.9 | 0.801 | 0.979 |
| Fn1 | Timp1 | 0 | 0.178 | 0 | 0.9 | 0.717 | 0.974 |
| Serpine1 | Timp1 | 0 | 0.172 | 0 | 0.9 | 0.713 | 0.974 |
| Fn1 | Vcan | 0 | 0.102 | 0 | 0.9 | 0.721 | 0.972 |
| Cd44 | Fn1 | 0 | 0.08 | 0.181 | 0.8 | 0.724 | 0.952 |
| Timp1 | Vcan | 0 | 0.122 | 0 | 0.9 | 0.498 | 0.952 |
| Fn1 | Tnc | 0.584 | 0.149 | 0.181 | 0.9 | 0.499 | 0.939 |
| Spp1 | Vcan | 0 | 0.063 | 0 | 0.9 | 0.371 | 0.935 |
| Tnc | Vcan | 0 | 0.166 | 0.174 | 0.9 | 0.157 | 0.934 |
| Itga5 | Spp1 | 0 | 0.063 | 0.148 | 0.9 | 0.199 | 0.927 |
| Itgav | Tnc | 0 | 0.063 | 0.212 | 0.9 | 0.064 | 0.921 |
| Timp1 | Tnc | 0 | 0.148 | 0 | 0.9 | 0.14 | 0.92 |
| Fn1 | Itgav | 0 | 0.064 | 0.801 | 0 | 0.6 | 0.919 |
| Penk | Timp1 | 0 | 0 | 0 | 0.9 | 0.144 | 0.91 |
| Spp1 | Tnc | 0 | 0.063 | 0 | 0.9 | 0.112 | 0.909 |
| Fn1 | Penk | 0 | 0 | 0 | 0.9 | 0.064 | 0.902 |
| Penk | Vcan | 0 | 0.063 | 0 | 0.9 | 0 | 0.902 |
| Penk | Tnc | 0 | 0.063 | 0 | 0.9 | 0 | 0.902 |
| Penk | Spp1 | 0 | 0 | 0 | 0.9 | 0.059 | 0.901 |
| Col4a1 | Itga5 | 0 | 0.082 | 0.067 | 0.6 | 0.336 | 0.742 |
| Col4a1 | Fn1 | 0 | 0.323 | 0.152 | 0 | 0.552 | 0.72 |
| Cd44 | Vcan | 0 | 0.066 | 0.181 | 0 | 0.646 | 0.706 |
| Kif22 | Prc1 | 0 | 0.554 | 0.18 | 0 | 0.235 | 0.695 |
| Anln | Prc1 | 0 | 0.585 | 0 | 0 | 0.274 | 0.686 |
| Itga5 | Tnc | 0 | 0.063 | 0.212 | 0.6 | 0.057 | 0.684 |
| Anlnl1 | Prc1 | 0 | 0.576 | 0 | 0 | 0.274 | 0.679 |
| Cd44 | Col4a1 | 0 | 0.061 | 0.09 | 0.6 | 0.121 | 0.659 |
| Cd44 | Timp1 | 0 | 0.223 | 0 | 0 | 0.577 | 0.658 |
| Cenpn | Prc1 | 0 | 0.617 | 0 | 0 | 0 | 0.617 |
| Cd44 | Itgav | 0 | 0.063 | 0.108 | 0 | 0.561 | 0.601 |
| Fn1 | S100a4 | 0 | 0.117 | 0 | 0 | 0.561 | 0.596 |
| Kpna2 | Prc1 | 0 | 0.58 | 0 | 0 | 0 | 0.58 |
| Serpine1 | Tnfrsf12a | 0 | 0.482 | 0 | 0 | 0.222 | 0.58 |
| Timp1 | Tnfrsf12a | 0 | 0.495 | 0 | 0 | 0.2 | 0.579 |
| Serpine1 | Spp1 | 0 | 0.09 | 0 | 0 | 0.542 | 0.565 |
| Cenpn | Kif22 | 0 | 0.551 | 0 | 0 | 0 | 0.551 |
| S100a4 | Timp1 | 0 | 0.366 | 0 | 0 | 0.302 | 0.539 |
| Itgav | Spp1 | 0 | 0.063 | 0.181 | 0 | 0.417 | 0.514 |
| Plp2 | Timp1 | 0 | 0.512 | 0 | 0 | 0 | 0.512 |
| Cd44 | Itga5 | 0 | 0.064 | 0.108 | 0 | 0.459 | 0.509 |
| Prc1 | Tubb4b | 0 | 0.064 | 0.396 | 0 | 0.175 | 0.492 |
| Praf2 | S100a4 | 0 | 0.063 | 0 | 0 | 0.47 | 0.482 |
| Col4a1 | Timp1 | 0 | 0.149 | 0.086 | 0 | 0.371 | 0.468 |
| Cd44 | S100a4 | 0 | 0.203 | 0 | 0 | 0.321 | 0.436 |
| S100a4 | Spp1 | 0 | 0.13 | 0 | 0 | 0.345 | 0.406 |
| Plp2 | S100a4 | 0 | 0.402 | 0 | 0 | 0 | 0.402 |
| Cenpn | Kpna2 | 0 | 0.4 | 0 | 0 | 0 | 0.4 |

## References

Brott, D., Gould, S., Jones, H., Schofield, J., Prior, H., Valentin, J.P., Bjurstrom, S., Kenne, K., Schuppe-Koistinen, I., Katein, A., Foster-Brown, L., Betton, G., Richardson, R., Evans, G., Louden, C., 2005. Biomarkers of drug-induced vascular injury, in: Toxicology and Applied Pharmacology. Academic Press Inc., pp. 441–445. https://doi.org/10.1016/j.taap.2005.04.028

Chen, D., Wang, X., Liang, D., Gordon, J., Mittal, A., Manley, N., Degenhardt, K., Astrof, S., 2015. Fibronectin signals through integrin α5β1 to regulate cardiovascular development in a cell type-specific manner. Developmental Biology 407, 195–210. https://doi.org/10.1016/j.ydbio.2015.09.016

Daguès, N., Pawlowski, V., Guigon, G., Ledieu, D., Sobry, C., Hanton, G., Freslon, J.L., Chevalier, S., 2007a. Altered gene expression in rat mesenteric tissue following in vivo exposure to a phosphodiesterase 4 inhibitor. Toxicology and Applied Pharmacology 218, 52–63. https://doi.org/10.1016/j.taap.2006.10.018

Daguès, N., Pawlowski, V., Sobry, C., Hanton, G., Borde, F., Soler, S., Freslon, J.L., Chevalier, S., 2007b. Investigation of the molecular mechanisms preceding PDE4 inhibitor-induced vasculopathy in rats: Tissue inhibitor of metalloproteinase 1, a potential predictive biomarker. Toxicological Sciences 100, 238–247. https://doi.org/10.1093/toxsci/kfm161

Dalmas, D.A., Scicchitano, M.S., Chen, Y., Kane, J., Mirabile, R., Schwartz, L.W., Thomas, H.C., Boyce, R.W., 2008. Transcriptional Profiling of Laser Capture Microdissected Rat Arterial Elements: Fenoldopam-induced Vascular Toxicity as a Model System. Toxicologic Pathology 36, 496–519. https://doi.org/10.1177/0192623307311400

Dalmas, D.A., Scicchitano, M.S., Mullins, D., Hughes-Earle, A., Tatsuoka, K., Magid-Slav, M., Frazier, K.S., Thomas, H.C., 2011. Potential candidate genomic biomarkers of drug induced vascular injury in the rat. Toxicology and Applied Pharmacology 257, 284–300. https://doi.org/10.1016/j.taap.2011.09.015

Edgar, R., Domrachev, M., Lash, A.E., 2002. Gene Expression Omnibus: NCBI gene expression and hybridization array data repository. Nucleic Acids Research 30, 207–210. https://doi.org/10.1093/nar/30.1.207

Fu, B.M., Tarbell, J.M., 2013. Mechano-sensing and transduction by endothelial surface glycocalyx: Composition, structure, and function. Wiley Interdisciplinary Reviews: Systems Biology and Medicine 5, 381–390. https://doi.org/10.1002/wsbm.1211

Gautier, L., Cope, L., Bolstad, B.M., Irizarry, R.A., 2004. Affy - Analysis of Affymetrix GeneChip data at the probe level. Bioinformatics 20, 307–315. https://doi.org/10.1093/bioinformatics/btg405

Heydarkhan-Hagvall, S., Chien, S., Nelander, S., Li, Y.C., Yuan, S., Lao, J., Haga, J.H., Lian, I., Nguyen, P., Risberg, B., Li, Y.S., 2006. DNA microarray study on gene expression profiles in co-cultured endothelial and smooth muscle cells in response to 4-and 24-h shear stress. Molecular and Cellular Biochemistry 281, 1–15. https://doi.org/10.1007/s11010-006-0168-6

Imanaka-Yoshida, K., Yoshida, T., Miyagawa-Tomita, S., 2014. Tenascin-C in development and disease of blood vessels. Anatomical Record 297, 1747–1757. https://doi.org/10.1002/ar.22985

Johansson, S., 1981. Cardiovascular lesions in Sprague-Dawley rats induced by long-term treatment with caffeine. Acta Pathol Microbiol Scand A 89, 185–91.

Kauffmann, A., Gentleman, R., Huber, W., 2009. arrayQualityMetrics - A bioconductor package for quality assessment of microarray data. Bioinformatics 25, 415–416. https://doi.org/10.1093/bioinformatics/btn647

Kavanaugh, A., Mease, P.J., Gomez-Reino, J.J., Adebajo, A.O., Wollenhaupt, J., Gladman, D.D., Lespessailles, E., Hall, S., Hochfeld, M., Hu, C., Hough, D., Stevens, R.M., Schett, G., 2014. Treatment of psoriatic arthritis in a phase 3 randomised, placebo-controlled trial with apremilast, an oral phosphodiesterase 4 inhibitor. Annals of the Rheumatic Diseases 73, 1020–1026. https://doi.org/10.1136/annrheumdis-2013-205056

Kerns, W., Schwartz, L., Blanchard, K., Burchiel, S., Essayan, D., Fung, E., Johnson, R., Lawton, M., Louden, C., MacGregor, J., Miller, F., Nagarkatti, P., Robertson, D., Sistare, F., Snyder, P., Thomas, H., Wagner, B., Ward, A., Jun, Z., 2005. Drug-induced vascular injury - A quest for biomarkers. Toxicology and Applied Pharmacology 203, 62–87. https://doi.org/10.1016/j.taap.2004.08.001

Kovacic, J.C., Dimmeler, S., Harvey, R.P., Finkel, T., Aikawa, E., Krenning, G., Baker, A.H., 2019. Endothelial to Mesenchymal Transition in Cardiovascular Disease: JACC State-of-the-Art Review. J Am Coll Cardiol. https://doi.org/10.1016/j.jacc.2018.09.089

Kuhn, M., Vaughan, D., 2020. yardstick: Tidy Characterizations of Model Performance.

Leek, J.T., Johnson, W.E., Parker, H.S., Jaffe, A.E., Storey, J.D., 2012. The SVA package for removing batch effects and other unwanted variation in high-throughput experiments. Bioinformatics 28, 882–883. https://doi.org/10.1093/bioinformatics/bts034

Liberzon, A., Birger, C., Thorvaldsdóttir, H., Ghandi, M., Mesirov, J.P., Tamayo, P., 2015. The Molecular Signatures Database Hallmark Gene Set Collection. Cell Systems 1, 417–425. https://doi.org/10.1016/j.cels.2015.12.004

Liberzon, A., Subramanian, A., Pinchback, R., Thorvaldsdóttir, H., Tamayo, P., Mesirov, J.P., 2011. Molecular signatures database (MSigDB) 3.0. Bioinformatics 27, 1739–1740. https://doi.org/10.1093/bioinformatics/btr260

Lok, Z.S.Y., Lyle, A.N., 2019. Osteopontin in Vascular Disease. Arterioscler Thromb Vasc Biol 39, 613–622. https://doi.org/10.1161/ATVBAHA.118.311577

Louden, C., Brott, D., Katein, A., Kelly, T., Gould, S., Jones, H., Betton, G., Valetin, J.P., Richardson, R., 2006a. Biomarkers and mechanisms of drug-induced vascular injury in non-rodents. Toxicologic Pathology. https://doi.org/10.1080/01926230500512076

Louden, C., Brott, D., Katein, A., Kelly, T., Gould, S., Jones, H., Betton, G., Valetin, J.P., Richardson, R., 2006b. Biomarkers and mechanisms of drug-induced vascular injury in non-rodents. Toxicologic Pathology. https://doi.org/10.1080/01926230500512076

Mikaelian, I., Cameron, M., Dalmas, D.A., Enerson, B.E., Gonzalez, R.J., Guionaud, S., Hoffmann, P.K., King, N.M.P., Lawton, M.P., Scicchitano, M.S., Smith, H.W., Thomas, R.A., Weaver, J.L., Zabka, T.S., Morton, D., Tomlinson, L., 2014. Nonclinical Safety Biomarkers of Drug-induced Vascular Injury:Current Status and Blueprint for the Future. Toxicologic Pathology 42, 635–657. https://doi.org/10.1177/0192623314525686

Morton, D., Houle, C.D., 2014. Perspectives on Drug-induced Vascular Injury. Toxicologic Pathology 42, 633–634. https://doi.org/10.1177/0192623314524490

Naba, A., Clauser, K.R., Hoersch, S., Liu, H., Carr, S.A., Hynes, R.O., 2012. The matrisome: in silico definition and in vivo characterization by proteomics of normal and tumor extracellular matrices. Mol Cell Proteomics 11, M111.014647. https://doi.org/10.1074/mcp.M111.014647

Pearson, T.A., Mensah, G.A., Alexander, R.W., Anderson, J.L., Cannon, R.O., Criqui, M., Fadl, Y.Y., Fortmann, S.P., Hong, Y., Myers, G.L., Rifai, N., Smith, S.C., Taubert, K., Tracy, R.P., Vinicor, F., 2003. Markers of inflammation and cardiovascular disease: Application to clinical and public health practice: A statement for healthcare professionals from the centers for disease control and prevention and the American Heart Association. Circulation 107, 499–511. https://doi.org/10.1161/01.CIR.0000052939.59093.45

Ponticos, M., Smith, B.D., 2014. Extracellular matrix synthesis in vascular disease: hypertension, and atherosclerosis. Journal of Biomedical Research 28, 25. https://doi.org/10.7555/JBR.27.20130064

Ritchie, M.E., Phipson, B., Wu, D., Hu, Y., Law, C.W., Shi, W., Smyth, G.K., 2015. Limma powers differential expression analyses for RNA-sequencing and microarray studies. Nucleic Acids Research 43, e47. https://doi.org/10.1093/nar/gkv007

Shimoyama, M., De Pons, J., Hayman, G.T., Laulederkind, S.J.F., Liu, W., Nigam, R., Petri, V., Smith, J.R., Tutaj, M., Wang, S.J., Worthey, E., Dwinell, M., Jacob, H., 2015. The Rat Genome Database 2015: Genomic, phenotypic and environmental variations and disease. Nucleic Acids Research 43, D743–D750. https://doi.org/10.1093/nar/gku1026

Slim, R.M., Yunling Song, Albassam, M., Dethloff, L.A., 2003. Apoptosis and Nitrative Stress Associated with Phosphodiesterase Inhibitor-Induced Mesenteric Vasculitis in Rats. Toxicologic Pathology 31, 638–645. https://doi.org/10.1080/01926230390241972

Smedley, D., Haider, S., Durinck, S., Pandini, L., Provero, P., Allen, J., Arnaiz, O., Awedh, M.H., Baldock, R., Barbiera, G., Bardou, P., Beck, T., Blake, A., Bonierbale, M., Brookes, A.J., Bucci, G., Buetti, I., Burge, S., Cabau, C., Carlson, J.W., Chelala, C., Chrysostomou, C., Cittaro, D., Collin, O., Cordova, R., Cutts, R.J., Dassi, E., Genova, A. Di, Djari, A., Esposito, A., Estrella, H., Eyras, E., Fernandez-Banet, J., Forbes, S., Free, R.C., Fujisawa, T., Gadaleta, E., Garcia-Manteiga, J.M., Goodstein, D., Gray, K., Guerra-Assunção, J.A., Haggarty, B., Han, D.-J., Han, B.W., Harris, T., Harshbarger, J., Hastings, R.K., Hayes, R.D., Hoede, C., Hu, S., Hu, Z.-L., Hutchins, L., Kan, Z., Kawaji, H., Keliet, A., Kerhornou, A., Kim, S., Kinsella, R., Klopp, C., Kong, L., Lawson, D., Lazarevic, D., Lee, J.-H., Letellier, T., Li, C.-Y., Lio, P., Liu, C.-J., Luo, J., Maass, A., Mariette, J., Maurel, T., Merella, S., Mohamed, A.M., Moreews, F., Nabihoudine, I., Ndegwa, N., Noirot, C., Perez-Llamas, C., Primig, M., Quattrone, A., Quesneville, H., Rambaldi, D., Reecy, J., Riba, M., Rosanoff, S., Saddiq, A.A., Salas, E., Sallou, O., Shepherd, R., Simon, R., Sperling, L., Spooner, W., Staines, D.M., Steinbach, D., Stone, K., Stupka, E., Teague, J.W., Dayem Ullah, A.Z., Wang, J., Ware, D., Wong-Erasmus, M., Youens-Clark, K., Zadissa, A., Zhang, S.-J., Kasprzyk, A., 2015. The BioMart community portal: an innovative alternative to large, centralized data repositories. Nucleic Acids Research 43, W589–W598. https://doi.org/10.1093/nar/gkv350

Sobota, J.T., 1989. Review of cardiovascular findings in humans treated with minoxidil. Toxicologic Pathology 17, 193–202. https://doi.org/10.1177/019262338901700115

Szklarczyk, D., Gable, A.L., Lyon, D., Junge, A., Wyder, S., Huerta-Cepas, J., Simonovic, M., Doncheva, N.T., Morris, J.H., Bork, P., Jensen, L.J., Von Mering, C., 2019. STRING v11: Protein-protein association networks with increased coverage, supporting functional discovery in genome-wide experimental datasets. Nucleic Acids Research 47, D607–D613. https://doi.org/10.1093/nar/gky1131

Tagliabracci, V.S., Wiley, S.E., Guo, X., Kinch, L.N., Durrant, E., Wen, J., Xiao, J., Cui, J., Nguyen, K.B., Engel, J.L., Coon, J.J., Grishin, N., Pinna, L.A., Pagliarini, D.J., Dixon, J.E., 2015. A Single Kinase Generates the Majority of the Secreted Phosphoproteome. Cell 161, 1619–1632. https://doi.org/10.1016/j.cell.2015.05.028

Weaver, J.L., Jun Zhang, Knapton, A., Miller, T., Espandiari, P., Smith, R., Gu, Y.Z., Snyder, R.D., 2010. Early events in vascular injury in the rat induced by the phosphodiesterase IV inhibitor SCH 351591. Toxicologic Pathology 38, 738–744. https://doi.org/10.1177/0192623310374331

Weaver, J.L., Snyder, R., Knapton, A., Herman, E.H., Honchel, R., Miller, T., Espandiari, P., Smith, R., Gu, Y.Z., Goodsaid, F.M., Rosenblum, I.Y., Sistare, F.D., Zhang, J., Hanig, J., 2008. Biomarkers in peripheral blood associated with vascular injury in Sprague-Dawley rats treated with the phosphodiesterase IV inhibitors SCH 351591 or SCH 534385. Toxicologic Pathology 36, 840–849. https://doi.org/10.1177/0192623308322310

Yu, G., Wang, L.G., Han, Y., He, Q.Y., 2012. ClusterProfiler: An R package for comparing biological themes among gene clusters. OMICS A Journal of Integrative Biology 16, 284–287. https://doi.org/10.1089/omi.2011.0118

Zhang, J., Herman, E.H., Robertson, D.G., Reily, M.D., Knapton, A., Ratajczak, H. v., Rifai, N., Honchel, R., Blanchard, K.T., Stoll, R.E., Sistare, F.D., 2006. Mechanisms and biomarkers of cardiovascular injury induced by phosphodiesterase inhibitor III SK&F 95654 in the spontaneously hypertensive rat. Toxicologic Pathology 34, 152–163. https://doi.org/10.1080/01926230600588562

Kuhn, M., Vaughan, D., 2020. yardstick: Tidy Characterizations of Model Performance.

Smedley, D., Haider, S., Durinck, S., Pandini, L., Provero, P., Allen, J., Arnaiz, O., Awedh, M.H., Baldock, R., Barbiera, G., Bardou, P., Beck, T., Blake, A., Bonierbale, M., Brookes, A.J., Bucci, G., Buetti, I., Burge, S., Cabau, C., Carlson, J.W., Chelala, C., Chrysostomou, C., Cittaro, D., Collin, O., Cordova, R., Cutts, R.J., Dassi, E. Genova, A. Di, Djari, A., Esposito, A., Estrella, H., Eyras, E., Fernandez-Banet, J., Forbes, S., Free, R.C., Fujisawa, T., Gadaleta, E., Garcia-Manteiga, J.M., Goodstein, D., Gray, K., Guerra-Assunção, J.A., Haggarty, B., Han, D.-J., Han, B.W., Harris, T., Harshbarger, J., Hastings, R.K., Hayes, R.D., Hoede, C., Hu, S., Hu, Z.-L., Hutchins, L., Kan, Z., Kawaji, H., Keliet, A., Kerhornou, A., Kim, S., Kinsella, R., Klopp, C., Kong, L., Lawson, D., Lazarevic, D., Lee, J.-H., Letellier, T., Li, C.-Y., Lio, P., Liu, C.-J., Luo, J., Maass, A., Mariette, J., Maurel, T., Merella, S., Mohamed, A.M., Moreews, F., Nabihoudine, I., Ndegwa, N., Noirot, C., Perez-Llamas, C., Primig, M., Quattrone, A., Quesneville, H., Rambaldi, D., Reecy, J., Riba, M., Rosanoff, S., Saddiq, A.A., Salas, E., Sallou, O., Shepherd, R., Simon, R., Sperling, L., Spooner, W., Staines, D.M., Steinbach, D., Stone, K., Stupka, E., Teague, J.W., Dayem Ullah, A.Z., Wang, J., Ware, D., Wong-Erasmus, M., Youens-Clark, K., Zadissa, A., Zhang, S.-J., Kasprzyk, A., 2015. The BioMart community portal: an innovative alternative to large, centralized data repositories. Nucleic Acids Research 43, W589–W598. https://doi.org/10.1093/nar/gkv350

