## Supplementary figures and images for "Identification of potential biomarker candidates of drug-induced vascular injury (DIVI) in rats using gene expression and histopathology data"

### S1_Figure

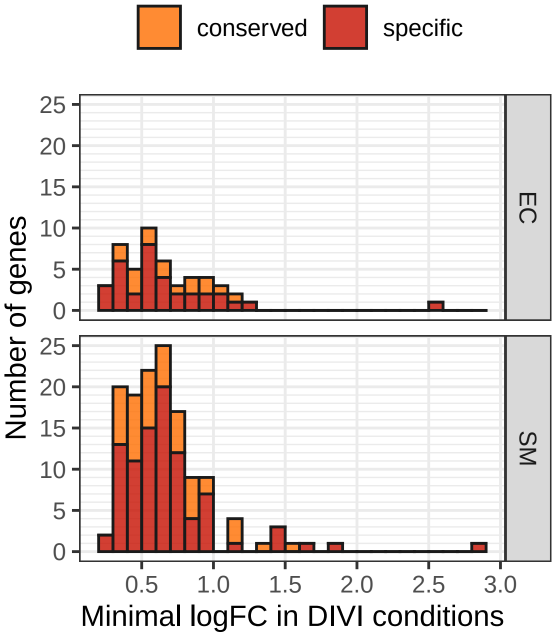

### S2_Figure

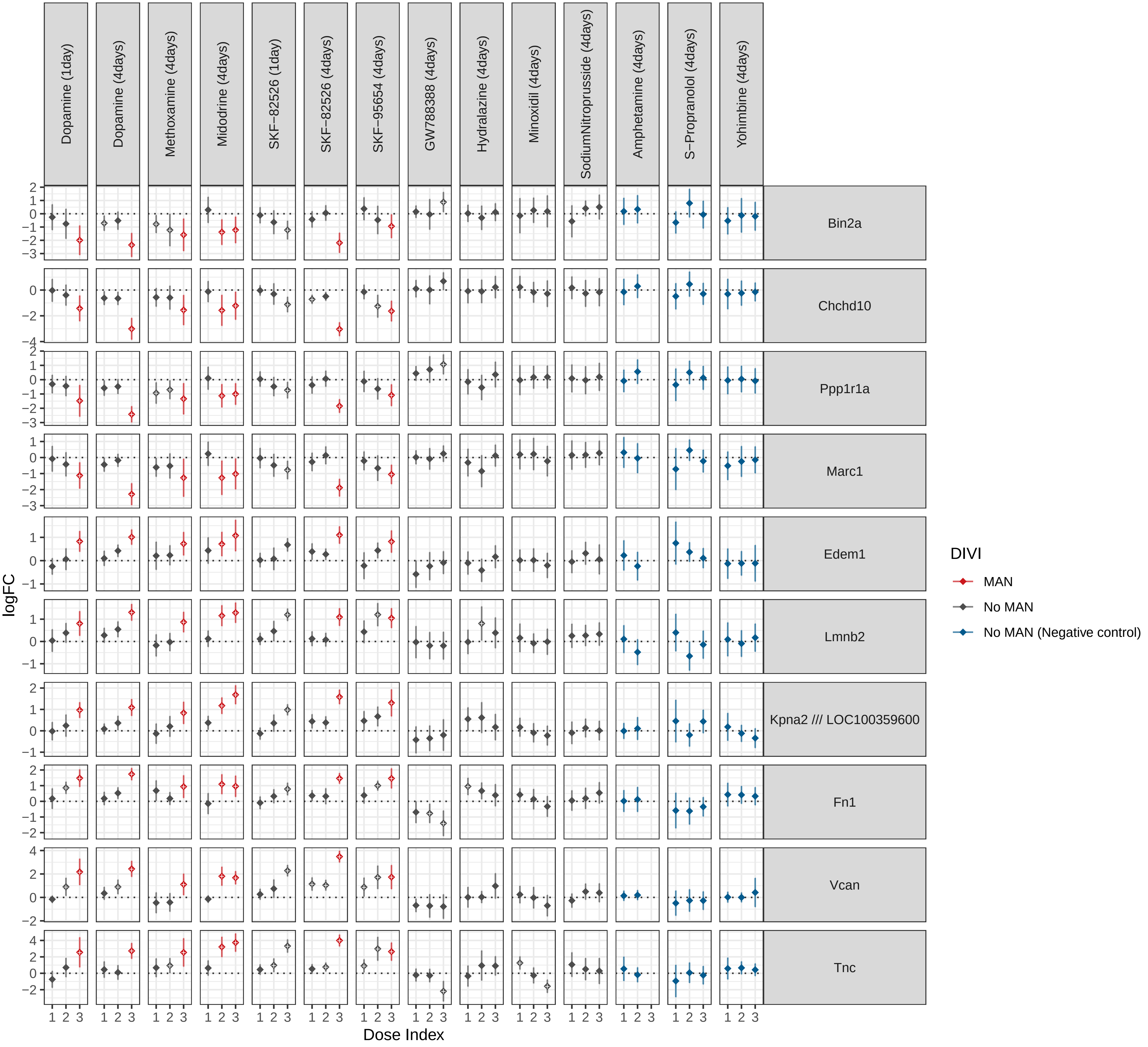

### S3_Figure

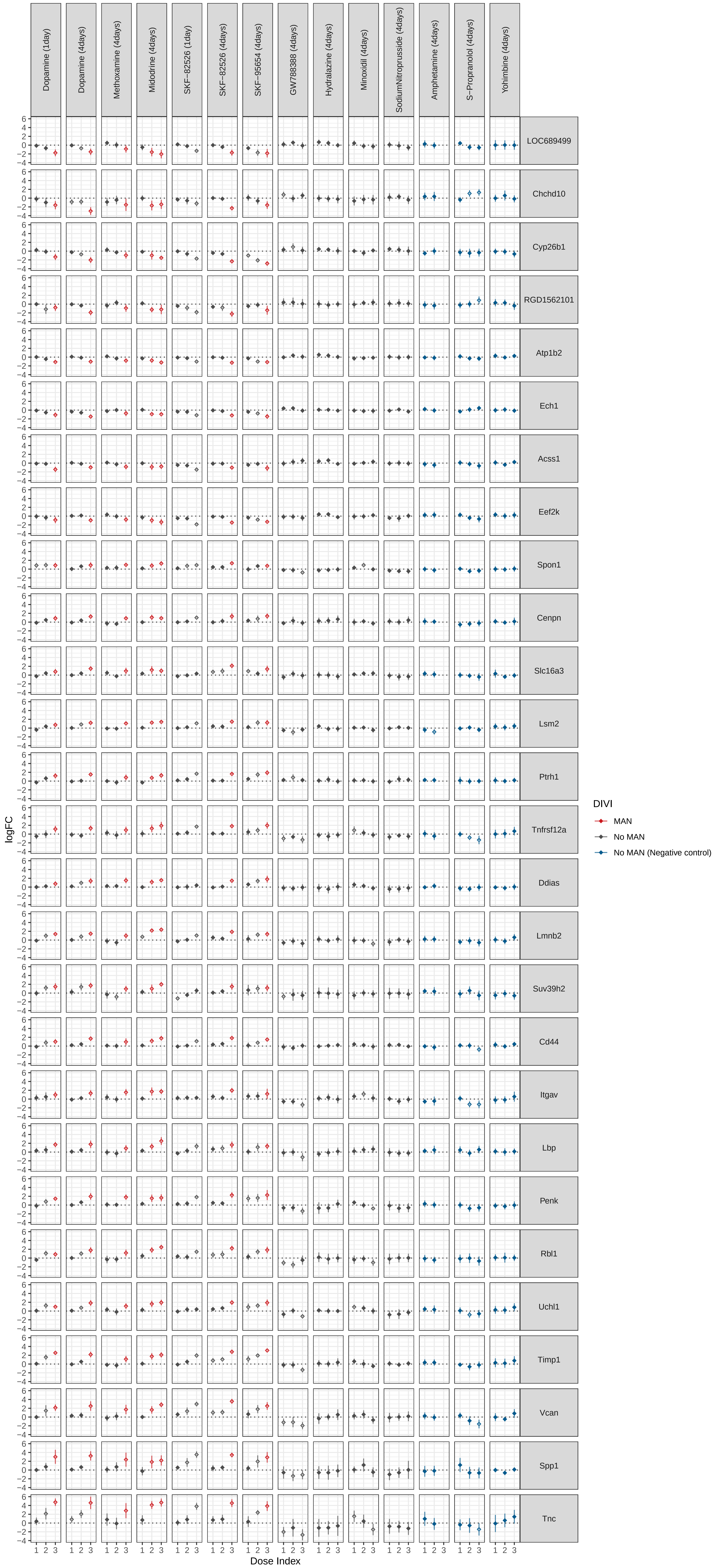

### S4_Figure

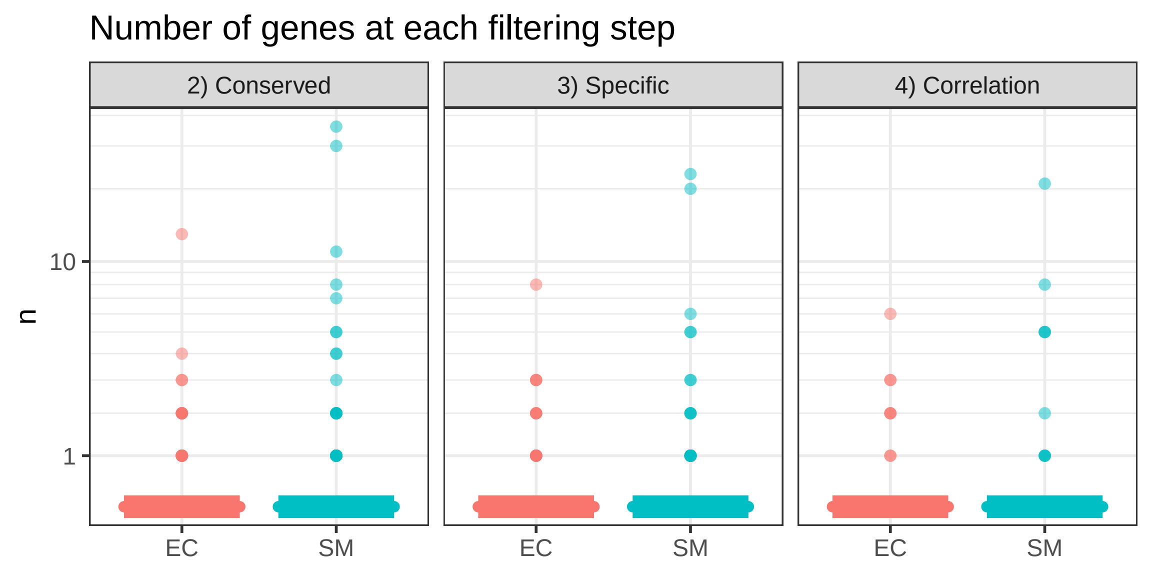

### S5_Figure

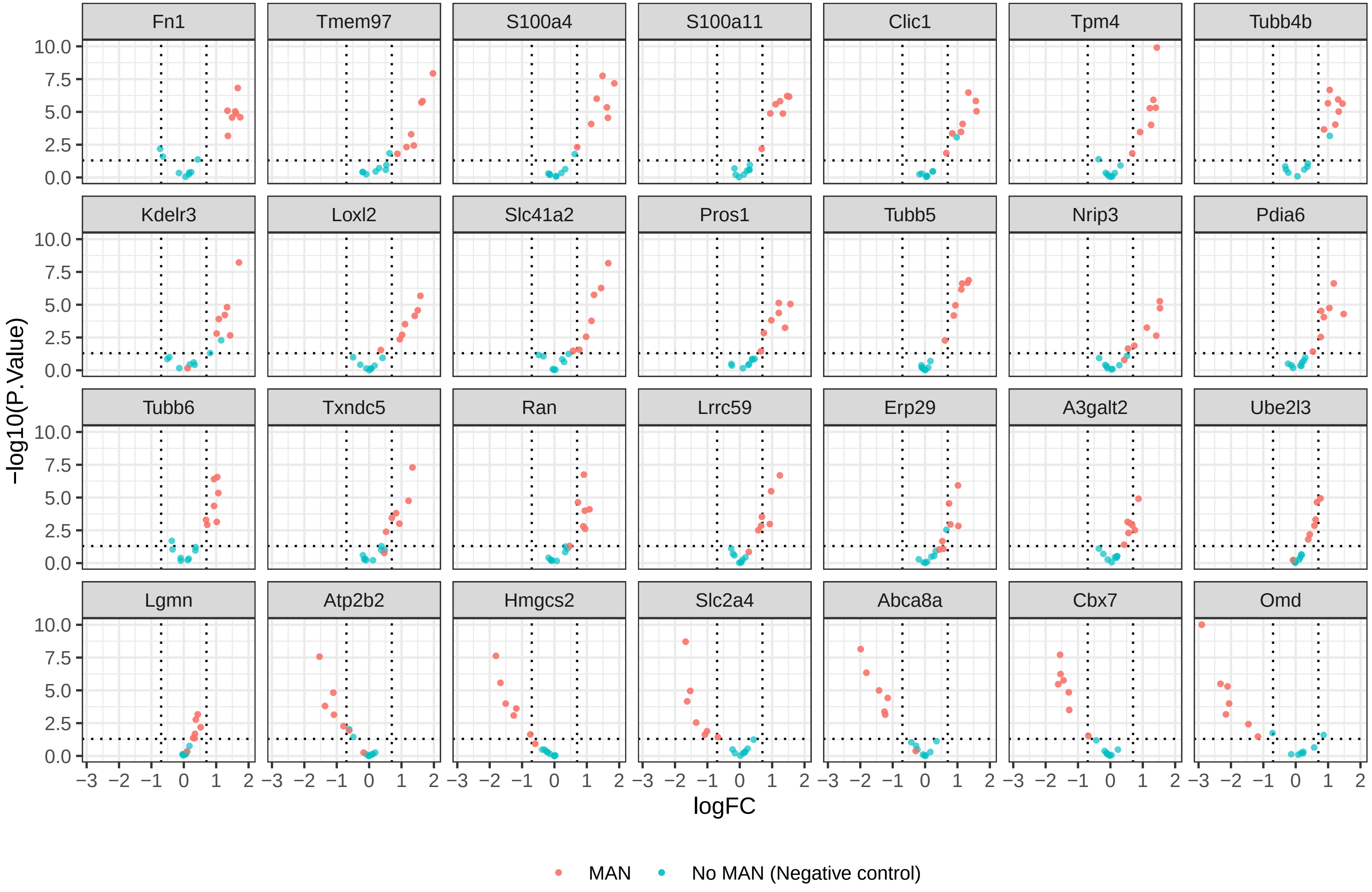

### S6_Figure

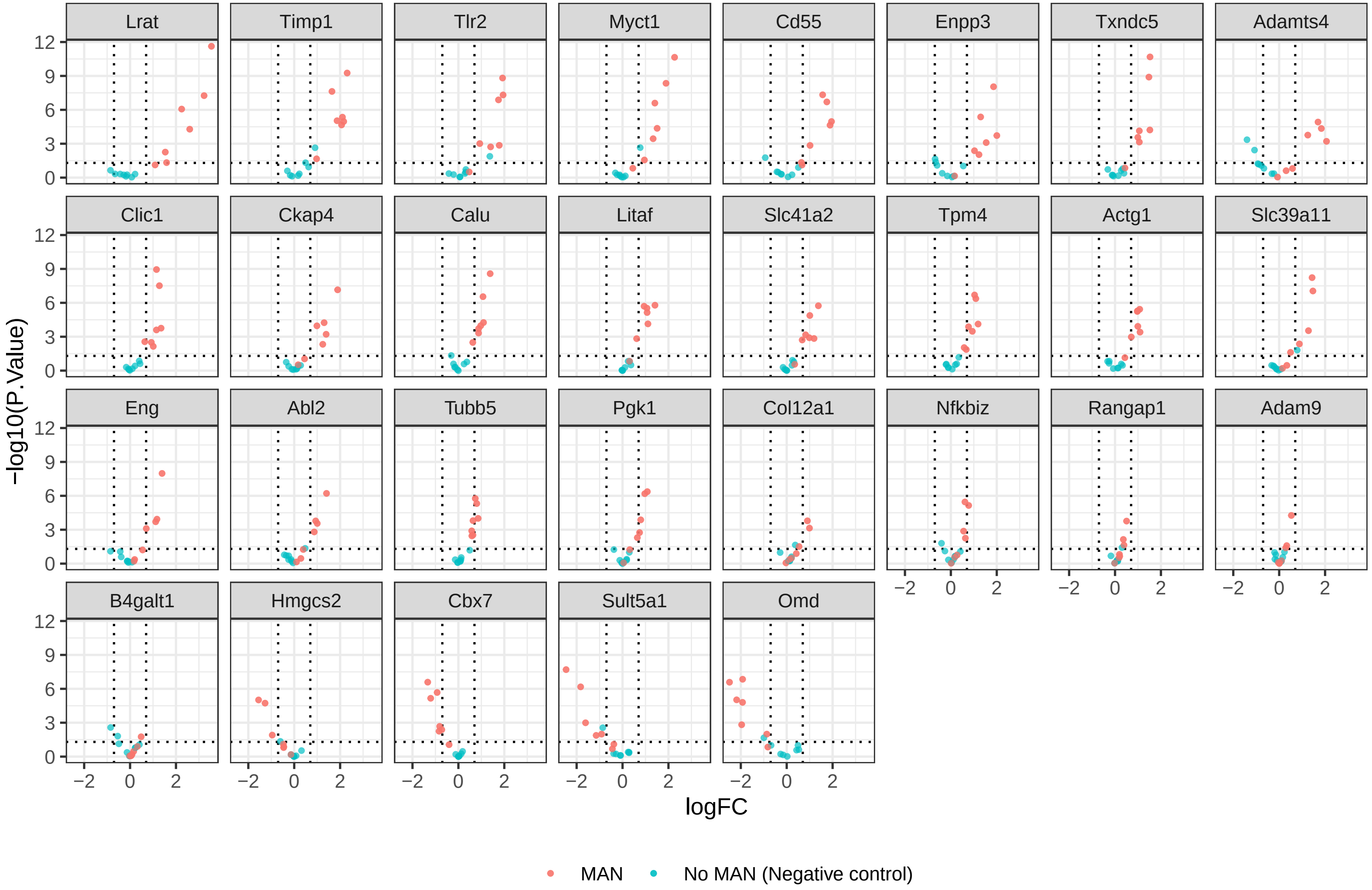
